# Immune checkpoint blockade induces gut microbiota translocation that augments extraintestinal anti-tumor immunity

**DOI:** 10.1101/2022.01.26.477865

**Authors:** Yongbin Choi, Yajing Gao, Laura A. Coughlin, Nicole Poulides, Jiwoong Kim, Xiaowei Zhan, Lora V. Hooper, Chandrashekhar Pasare, Andrew Y. Koh

## Abstract

Gut microbiota are critical for effective immune checkpoint blockade therapy (ICT) for cancer. The mechanisms by which gut microbiota augment extraintestinal anti-cancer immune responses, however, are largely unknown. Here, we find that ICT induces translocation of specific endogenous gut microbiota into secondary lymphoid organs and subcutaneous melanoma tumors. Mechanistically, gut microbiota activated dendritic cells (DCs) traffic a selective subset of gut bacteria to mesenteric lymph nodes (MLN) and promote optimal anti-tumor T-cell responses in both the tumor draining lymph nodes (TDLN) and the primary tumor. Antibiotic treatment resulted in decreased gut microbiota translocation into MLN and TDLN, diminished polyfunctional effector CD8+ T cell responses, and attenuated response to ICT. Our findings illuminate a key mechanism by which gut microbiota promote extraintestinal anti-cancer immunity.

**One sentence summary:** Following immune checkpoint blockade therapy, dendritic cells traffic gut microbiota into secondary lymphoid organs, promoting optimal extraintestinal anti-cancer immunity.

---

Immune checkpoint inhibitor therapy (ICT), targeting cytotoxic T-lymphocyte–associated-antigen-4 (CTLA-4) and/or programmed cell death protein 1 (PD-1), results in durable remissions in patients with previously incurable cancers (*1*). Yet up to 50% of cancer patients remain unresponsive to ICT (*2*). A variety of host associated factors have been implicated as a potential cause of this therapeutic discrepancy, and one of the most intriguing is the gut microbiome. There is growing evidence that the gut microbiome plays an instructive role in modulating cancer immunotherapy (*3-14*). Mice that lack gut microbiota or those pre-treated with antibiotics have a significantly diminished response to immune checkpoint inhibitor therapy (ICT) (*13, 14*). Our group (*7*) and others (*4, 8, 12*) have identified specific, yet distinct, gut microbiota associated with clinical response to cancer immunotherapy. And recent reports suggest that in cancer patients previously unresponsive to immunotherapy, fecal microbiota transplantation is both safe and potentially effective in augmenting anti-cancer immune responses (*15, 16*).

There is no consensus on exactly which gut microbiota are required for optimal host anti-cancer immune responses. A variety of gut microbiota species have been identified as correlated with positive clinical response to immunotherapy and/or effective in augmenting ICT responses in preclinical cancer models: *Bifidobacterium* spp. (*10, 13*), *Akkermansia muciniphila (12*), *Enterococcus* spp., (*5, 17*) and *Faecalibacterium prausnitzii (4, 7, 8*). To further confound matters, *Bacteroides* species have been shown to have both potentially beneficial (*7, 14*) and negative effects (*16, 18*) on cancer immunotherapy responses. Thus, it is unclear as to what generalizable “rules” can be ascertained from these data.

Gut microbiota can augment cancer immunotherapy responses through a variety of mechanisms. Bacteria have long been known to be potent activators of the innate immune system which can then prime T cells and induce anti-cancer immune responses (e.g. Coley’s Toxin) (*19*). Gut microbiota-derived metabolites (e.g. inosine (*9*) or short-chain fatty acids(*3*)) or pathogen-associated molecular patterns (e.g. muramyl peptide (*20*)) can enhance T cell anti-cancer responses. Finally, in some instances, specific endogenous gut microbiota may harbor antigens which cross-react with tumor antigens/neoantigens and thus via molecular mimicry increase T cell mediated anti-cancer responses (*21*).

Despite these recent intriguing findings, a major unanswered question is how do gut microbiota dictate or shape extraintestinal immune responses which promote tumor killing in distant sites. The potential immunologic influence or impact of gut microbiota on cancers that arise in the gastrointestinal tract (e.g. colorectal, pancreatic, and liver cancer) appears to be more obvious, given the close proximity of gut microbiota, immune cells, and the tumor. Yet, how gut microbiota modulate immune responses against extra-intestinal tumors, such as melanoma or lung cancer, is unclear.

Here, using a preclinical melanoma model and anti-PD-1/CTLA-4 therapy, we show that ICT induces translocation of specific gut microbiota into secondary lymphoid organs and tumor, thereby activating dendritic cells (DC) and priming anti-tumor T cell responses. Gut microbiota translocation into mesenteric lymph nodes (MLN) is essential for optimal anti-tumor T-cell responses in the tumor draining lymph node and tumor. DCs are critical not only for gut microbiota-dependent immune augmentation of ICT but also for facilitating gut microbiota translocation into MLN. Finally, antibiotic depletion of endogenous gut microbiota results in decreased gut microbiota translocation into MLN and TDLN, diminished effector T cell responses, and attenuated ICT responses. Together, our studies reveal a critical mechanism where gut microbiota translocation of specific immunogenic taxa, induced by ICT and aided by DCs, into secondary lymphoid organs results in optimal priming of anti-tumor responses effective against extraintestinal tumors.

## Immune checkpoint inhibitor therapy induces gut microbiota translocation into secondary lymphoid organs and tumor

We first sought to determine whether ICT, in the absence or presence of tumor, could induce gut microbiota translocation into secondary lymphoid organs. We administered ICT (combined anti-PD-1 and anti-CTLA-4 monoclonal antibody treatment) to mice with or without B16-F10 melanoma tumors and assessed bacterial translocation by culturing mesenteric lymph node (MLN) homogenates on various selective media under anaerobic conditions (fig. S1A). Indeed, mice receiving ICT had significantly greater number of cultured bacteria from MLNs (fig. S1B), including taxa previously reported to be associated with positive ICT response or implicated in host immune-anti cancer responses: *Bifidobacterium pseudolongum (13), Bacteroidetes thetaiotamicron (7, 22), Enterococcus faecalis (20, 22*), and *Lactobacillus spp*. (*22*) (fig. S1C). Interestingly, the presence of melanoma tumor alone did not result in a significant increase of cultured bacteria from MLNs, whereas the administration of ICT, regardless of tumor presence, resulted in a significant increase in the number of cultured bacteria in MLN tissue (fig. S1B).

We then asked whether there was a distinct temporal pattern of bacterial translocation throughout the course of ICT treatment. To address this question, we implanted B16-F10 tumors in a cohort of mice simultaneously and then proceeded with ICT treatment (fig. 1A). On each day, we selected mice (n=6-8) from different cages and harvested MLN, spleen, tumor draining lymph nodes (TDLN; defined as the right inguinal lymph node as tumors were implanted in the right flank), and tumor. Tissue microbiomes were assessed by both culturing of tissue homogenates and 16S rRNA sequencing of tissue gDNA. Notably, bacterial translocation was present but limited (< mean of 65 CFU/mg tissue) for all tissue types before the first dose of ICT (day 4) (fig. 1B). After initiation of ICT, bacterial translocation was readily detected in each tissue type (but to the greatest magnitude in MLN) and often increasing after each dose of ICT (fig. 1B). These data suggest that ICT induces gut microbiota translocation into secondary lymphoid tissues and tumor.

**Fig. 1.**
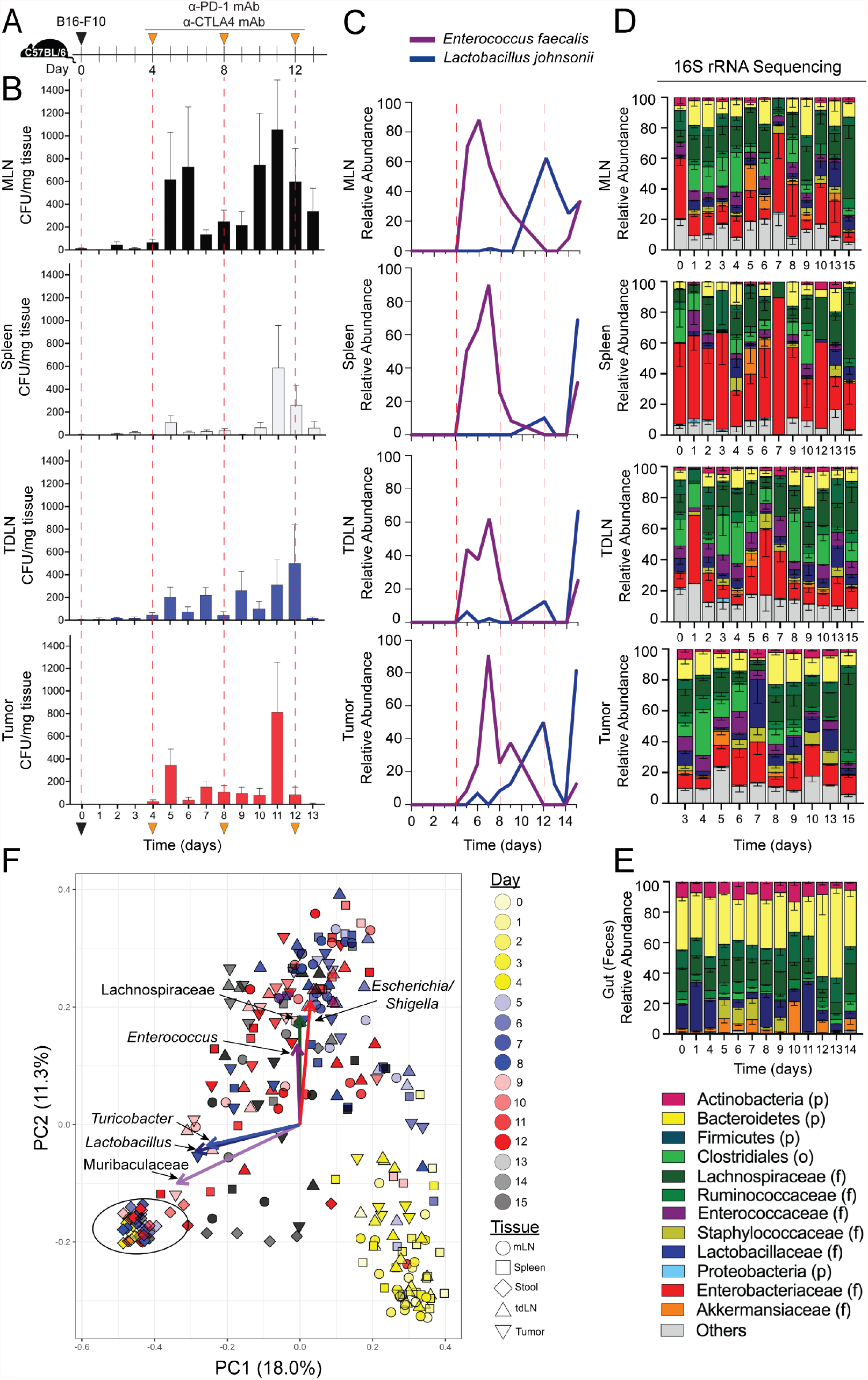
Immune Checkpoint Inhibitor Therapy (ICT) Induces Gut Microbiota Translocation into Secondary Lymphoid Organs and Tumor. (**A**) Schematic diagram of the strategy used to assess the temporal dynamics of bacterial translocation into secondary lymphoid organs and tumor in C57BL/6 mice (female, 6-8 wks, Jackson) bearing B16-F10 melanoma tumors and receiving ICT (anti-PD-1 and anti-CTLA-4 mAb, 200μg). (**B**) Cultured bacterial levels in MLN, Spleen, TDLN, and Tumor. Tissue homogenates were serially diluted, plated on BHI/Blood agar, and cultured at 37°C under anaerobic conditions for 24-72 hours. n=3-4 per time point per experiment. Two experiments were performed for a final n=6-8 per group. Bars represent the mean + SEM. (**C**) Relative abundance of cultured bacteria from secondary lymphoid organs and tumor, as determined by full length (V1-V9) 16S rRNA sequencing of gDNA isolated from cultured bacterial isolates. (**D-F**) 16sRNA gene sequencing (V4 region) of tissue gDNA isolated from mice as detailed in fig. 1A. (**D**) Relative abundance of microbiota in MLN, spleen, TDLN, as determined by 16S rRNA sequencing. (**E**) Relative abundance of microbiota in the gut (feces), as determined by 16S rRNA sequencing. (**F**) Principal coordinate analysis of tissue and gut 16S rRNA sequencing data, weighted and normalized by UniFrac distances. The proportion of variance accounted by each principal component is indicated. Vector analysis (as indicated by arrows) performed by singular value decomposition of 16S rRNA sequencing data interpreted visually as a linear biplot.

Upon taxonomic identification of cultured microbiota, two dominant taxa were noted: *Enterococcus faecalis* and *Lactobacillus johnsonii* (fig. 1C). A distinct temporal pattern of gut microbiota translocation was noted. *E. faecalis* was the most abundant gut microbiota translocator during the early phases of ICT treatment (between dose 1 and 2) while *L. johnsonii* exhibited dominance after the second dose of ICT. Of note, translocation of *E. faecalis* and *L. johnsonii* into secondary lymphoid organs has previously been reported as critical for augmenting anti-cancer immune responses in the setting of cyclophosphamide therapy (*22*) in preclinical models. Moreover, *E. faecalis* and *L. johnsonii* were recently identified as adept translocators into tumors in a preclinical colonic cancer model, with the latter showing efficacy in enhancing anti-CTLA-4 efficacy in this model (*9*). Finally, *L. johnsonii* and *Enterococcus* spp. have been associated with positive clinical response to ICT in melanoma patients (*15, 18*). Other notable cultured taxa include *Akkermansia muciniphila* (*12*) and *Bacteroides thetaiotaomicron* (*14*), which have been shown to augment anti-PD1 and anti-CTLA-4 responses respectively in preclinical models (fig S2).

Since the ability to successfully culture gut microbiota is highly variable, we sought to further characterize the secondary lymphoid organ and tumor microbiomes using 16S rRNA gene sequencing (fig. 1D and E). Using multidimensional scaling of the 16S rRNA data and factoring both tissue type and the time of sample collection, we noted that the gut microbiome taxonomic composition remained steady over the course of ICT (exhibited by the tight clustering of gut microbiome samples, diamonds, in the lower left-hand corner and demarcated with an ellipse, fig. 1F). In contrast, while lymphoid tissue and tumor samples collected prior to the first dose of ICT (days 0-4) also grouped together, regardless of tissue type (fig. 1F, lower right-hand corner, various tissue type samples all in yellow), notable taxonomic shifts occur after ICT initiation. By using singular value decomposition interpreted visually as a linear biplot (*23*), we identified six taxa as key drivers of taxonomic compositional changes in secondary lymphoid organs and tumor at later time points (days 5-15): *Enterococcus*, Lachnospiraceae, and *Escherichia/Shigella* accounting for the majority of change, with *Lactobacillus, Muribaculaceae*, and *Turibacter* contributing as well (fig 1F, demarcated as labeled arrows). When utilizing another well-established microbiome taxonomic differential abundance analysis tool (linear discriminant analysis effect size, LEfSe), we confirmed that *Enterococcus*, Lachnospiraceae (*Blautia* spp.) and *Shigella* were significantly enriched in tumor and secondary lymphoid organ tissues compared to the gut (fig. S3). Interestingly, microbiota taxa abundant in the gut (e.g. Bacteroidetes) were not necessarily highly abundant in the secondary lymphoid organs or tumors and *vice versa* (e.g. low abundance of Enterobacteriaceae in the gut, but high relative abundance in tissues) (fig. 1D and E). Finally, overall levels of bacteria (as determined by bacterial group qPCR of the same gDNA samples used for 16S rRNA sequencing shown in fig 1D, E) showed a comparable variation over time to that observed with the cultured microbiota abundance (fig 1B), with increases in bacterial levels after each ICT administration (fig. S4). Taken together, these results suggest that ICT induces translocation of specific gut microbiota, which are not necessarily abundant in the gut, into secondary lymphoid organs and tumor.

### Highly abundant microbiota translocators into secondary lymphoid organs activate DCs and induce anti-tumor effector T cell responses

We then asked whether the most abundant translocated bacteria identified by culturing and sequencing (specifically *Enterococcus* spp., *Lactobacillus johnsonii* and Enterobacteriaceae (fig. 1)) induce distinct immune responses to facilitate anti-cancer immunity. Translocated bacteria or bacterial components can provide a diverse array of pathogen-associated molecular patterns (PAMPs) capable of agonizing various pattern recognition receptors (PRRs) and thus activating innate immune responses (*24, 25*). Thus, we hypothesized that translocated gut bacterial components activate dendritic cells (DCs) that consequently drive tailored T cell immunity (*26*). Indeed, mouse DCs stimulated with *Enterococcus faecalis, Lactobacillus johnsonii*, and Enterobacteriaceae member *Escherichia coli* (fig. 2A) showed a significant increase in the surface expression of MHCII and co-stimulatory receptors, CD40, CD80 and CD86 (fig. 2B and C, fig. S5), whereas stimulation with *Lactobacillus acidophilus*, a common gut microbiota commensal and over-the-counter probiotic constituent, did not significantly activate DCs. We then investigated whether these gut microbiota translocators had a differential capacity to prime naïve T cells. We used an antigen-restricted DC-T cell priming system (e.g. MHC class-I-restricted, ovalbumin-specific, CD8+ T cells from OT-I TCR transgenic mice) and measured the activation and functional phenotypes of T cells after the co-culture with DCs (fig. 2D) (*26*). While OT-I CD8+ T cells primed with DCs pulsed with all bacteria showed significantly higher activation (fig. 2E), only DCs pulsed with highly abundant microbiota translocators *E. faecalis, L. johnsonii*, and *E. coli* induced significantly greater interferon-γ (IFN-γ) production in CD8+ T cells compared to the non-stimulated and *L. acidophilus* groups (fig. 2F). These data are consistent with prior published data emphasizing the importance of DCs in antitumor immunity in the setting of ICT (*27-29*), but additionally are concordant with our prior work highlighting the differential capacity of specific gut microbiota in driving specific T-cell immune responses (e.g. Th1 vs Th17, etc.) (*26*).

**Figure 2.**
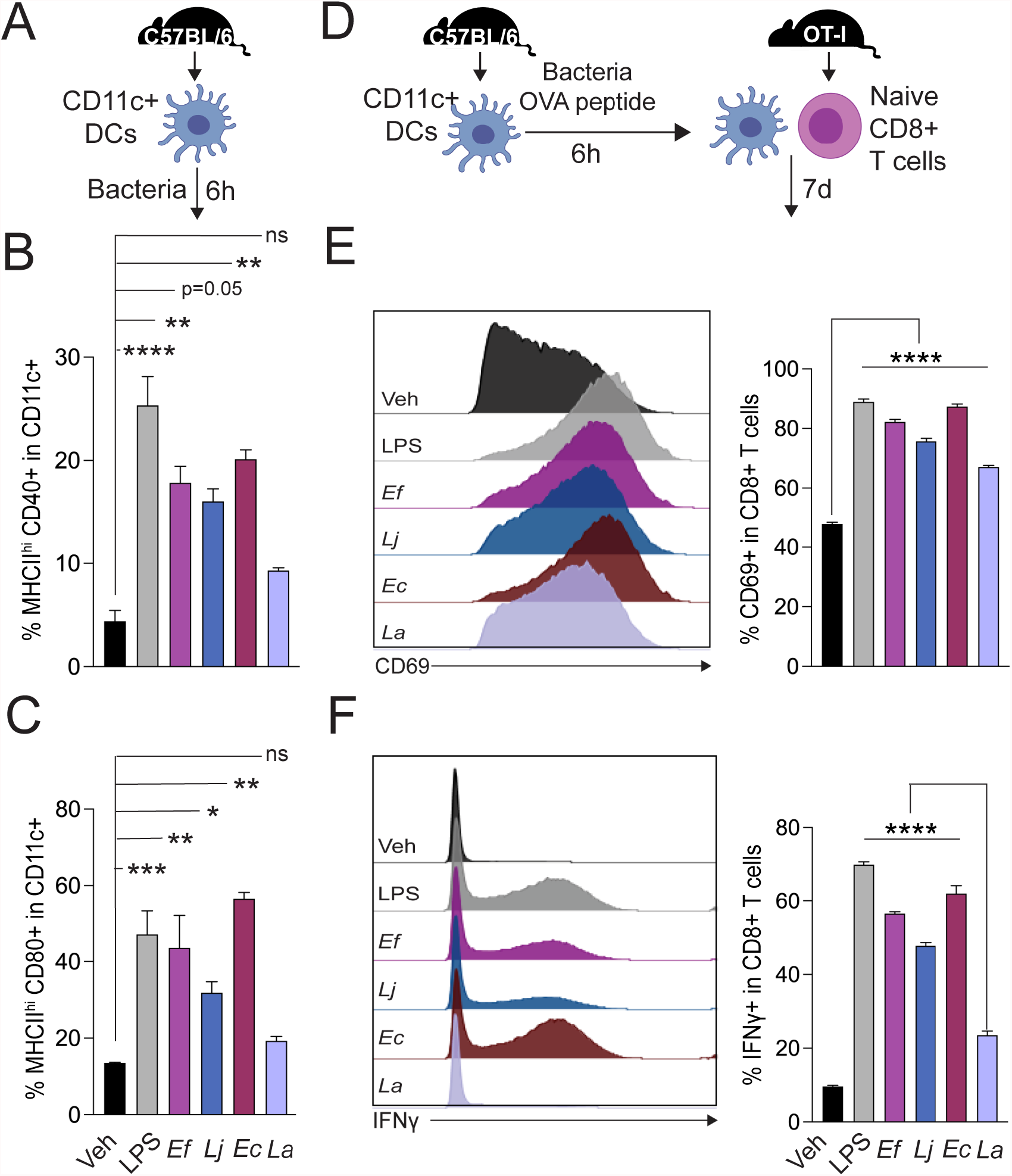
Highly abundant microbiota translocators into secondary lymphoid organs activate DCs and induce anti-tumor effector T cell responses. (**A**) Schematic overview of the protocol used to assess the dendritic cell (DC)-activating potential of various gut microbiota bacteria. CD11c+ DCs were isolated from the spleen of C57BL6/J mice (female, 6-8 wks, Jackson) bearing B16-FLT3L tumors. Isolated DCs were stimulated with vehicle (PBS), *Escherichia coli* LPS, *Enterococcus faecalis* (*Ef*), *Lactobacillus johnsonii* (*Lj*), *Escherichia coli* (*Ec*), or *Lactobacillus acidophilus* (La) for 6 hours. DCs were the analyzed by flow cytometry. Proportion of (**B**) MHC-II high, CD40+ cells and (**C**) MHC-II high, CD80+ cells among CD11c+ DCs. (**D**) Schematic overview of the protocol used to assess T cell activating and priming potential of DCs stimulated with different gut microbiota. CD11c+ DCs were pulsed with OVA 257-264 peptide and gut microbiota for 6h. Stimulated DCs were then co-cultured with naïve CD8+ T cells isolated from age-and sex-matched OT-I mice for 7d. Surface expression of T cell activation marker CD69 and intracellular interferon-γ (IFN-γ) were quantified by flow cytometry. (E) Proportion of (**E**) CD69+ and (**F**) IFN-γ+ cells among CD8+ T cells. Bars represent the mean + SEM. All assays were performed in triplicate. Statistical analysis by Mann-Whitney test. *P<0.05, **P<0.01, ***P<0.0.001, ****P<0.0001.

In an effort to ascertain whether there was any human clinical data to corroborate our murine findings, we interrogated tumor RNASeq data acquired from 458 human melanoma patients in the Cancer Genome Atlas. mRNA expression of toll-like receptors (TLR) 2, which senses bacterial structural components such as lipoteichoic acid found in gram-positive bacteria such as *L. johnsonii*, and TLR4, which senses the bacterial gram-negative component lipopolysaccharide found in bacteria such as *E. coli*, in melanoma was significantly associated with prolonged survival (fig. S6, Table S1). Further, the expression of CD40, CD80 and CD86 (which can be expressed by antigen presenting cells) was also significantly associated with patient survival (fig. S6, Table S1). Thus, while these data are correlative, it does beg the question as to whether bacterial structural component activation of antigen presenting cells could potentially influence survival outcomes in melanoma patients. Collectively, these results suggest that in order to achieve optimal anti-tumor immune responses, gut microbiota must not only translocate into secondary lymphoid organs and/or tumor but also be sufficiently immunogenic for the innate immune system to drive anti-cancer immune responses.

### Mesenteric lymph nodes modulate gut microbiota-dependent anti-tumor priming responses in the tumor-draining lymph node and tumor

We hypothesized that gut microbiota translocation into secondary lymphoid organs, particularly the mesenteric lymph nodes (MLN), was critical for shaping extra-intestinal anti-tumor responses in the tumor draining lymph node (TDLN) and the tumor. As a first step, we implanted melanoma tumors into lymphotoxin-α knockout mice (LTA KO) that lack secondary lymphoid organs, including MLN and inguinal lymph nodes, and Peyer’s patches (*30*) and then administered ICT. Indeed, LTA KO mice had a diminished response to ICT, despite having intact gut microbiota, compared to co-housed wild-type controls (fig. S7). Of note, LTA KO mice have normal quantitative and qualitative immune function (with the exception of a four-fold increase in B cells (*30*)) but a notable difference in gut microbiome composition (*31*) compared to wild-type mice. To further investigate the importance of secondary lymphoid organs in modulating gut microbiota dependent anti-tumor immune responses, we administered FTY-720 (fingolimod), a sphingosine-1-phosphate receptor modulator which sequesters lymphocytes in lymph nodes, to mice bearing melanoma tumors and receiving ICT. Mice receiving FTY-720 had larger tumor volumes compared to untreated controls, despite having intact gut microbiota and intact lymph nodes (fig. S8). These data suggest that secondary lymphoid organs and lymphocyte egress from these lymphoid tissues are critical for optimal gut microbiota-induced anti-cancer responses in the setting of ICT.

We then asked whether specific secondary lymphoid organs had a differential capacity to modulate gut microbiota-dependent extra-intestinal anti-tumor responses. Hence, we surgically resected different secondary lymphoid organs (MLN, TDLN, or spleen) from mice bearing melanoma tumors with intact gut microbiota and then administered ICT (fig. 3A). Interestingly, while all surgical groups exhibited larger tumor growth, only mice with MLN resection had significantly larger tumor volumes compared to control mice (receiving sham surgery) (fig. 3B). Similarly, while all surgical groups exhibited decreased survival time, only mice lacking MLN had significantly decreased survival compared to sham surgery control mice (p= 0.025, log-rank test, fig 3C). Further, MLN removal led to a significant decrease in bacterial load in TDLN and tumor, suggesting that MLN may play a role as a conduit of intact bacteria or gut microbiota-derived components from the gut to extra-intestinal sites, such as TDLN and/or tumor (fig. 3D and E).

**Figure 3.**
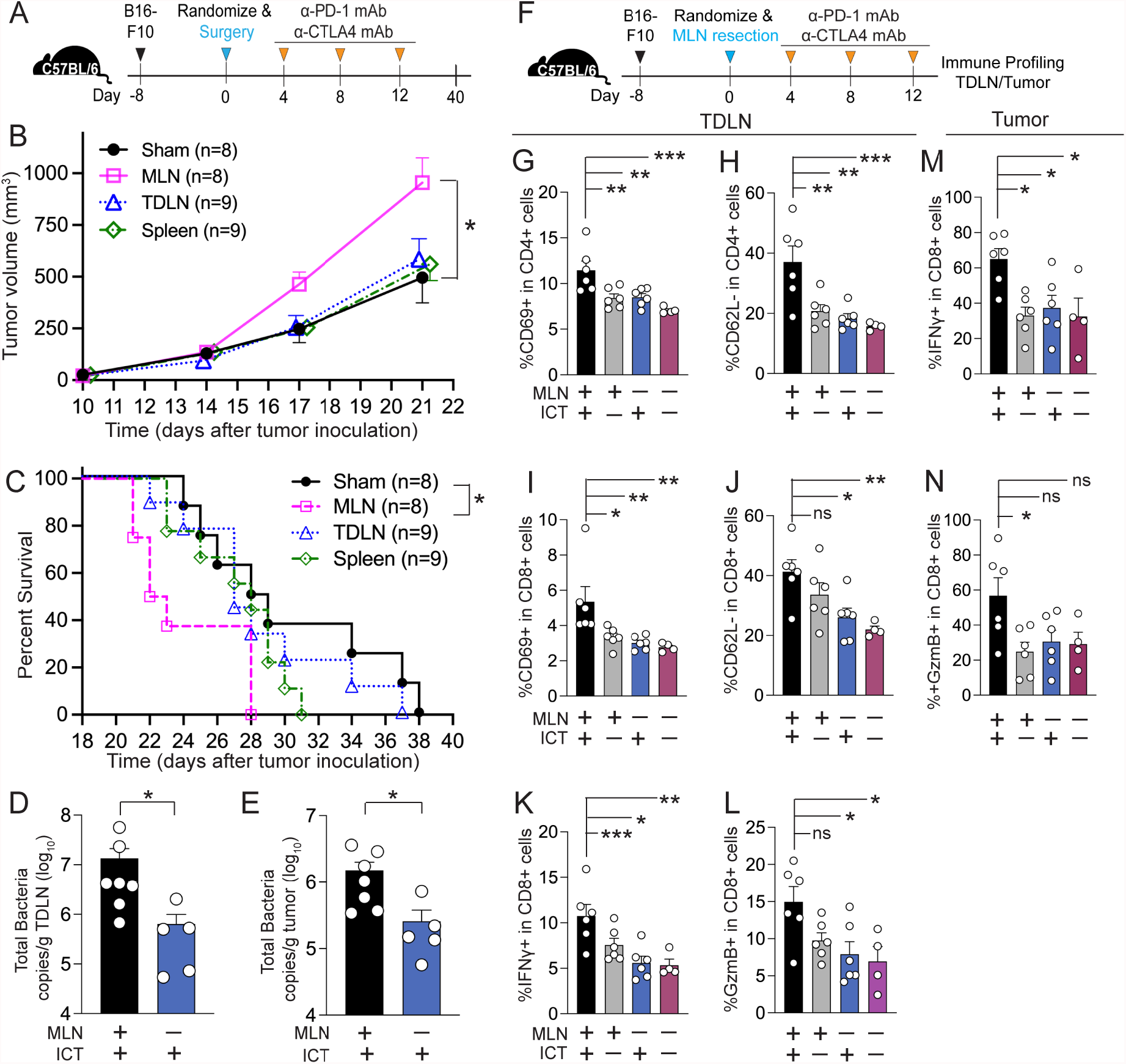
Mesenteric lymph nodes modulate gut microbiota-dependent anti-tumor priming responses in the tumor-draining lymph node and tumor. (**A**) Schematic overview of the protocol used to assess the impact of surgical resection of secondary lymphoid organs or control (sham surgery involving longitudinal abdominal incision only) on ICT efficacy. C57BL/6 mice (female, 6-8 wks, Jackson) were inoculated with 1 × 10^5^ B16-F10 cells subcutaneously in the right flank. Mice with comparable tumor volumes were randomized to receive surgery. 200μg anti-PD-1 and 200μg anti-CTLA-4 mAb (ICT) were injected intraperitoneally on days 4, 8, and 12 post-surgery. n=8-9 per group. (**B**) Tumor volume and (**C**) survival of tumor-bearing mice as detailed in fig 1A. Total bacterial load in tumor-draining lymph node (TDLN) (**D**) and tumor (**E**) of mice + MLN + ICT, as determined by bacterial group (Eubacteria, all bacteria) quantitative-PCR (qPCR) of tissue gDNA collected from mice as detailed in fig. 2A. (**F**) Schematic overview of the protocol used to assess the impact of MLN removal on the CD4+ and CD8+ T cell immune responses in TDLN and tumor of B16-F10 tumor bearing mice receiving ICT. n=4-6 per group (**G-L**) Quantification of CD4+ and CD8+ T cell immune response in the TDLN of mice + MLN by flow cytometry. The proportion of (**G, I**) activated (CD69+) and (**H, J**) effector (CD62L-) T cells among CD4+ T cells and CD8+ T cells respectively. The proportion of (**K**) IFN-γ and (**L**) granzyme B (GzmB) producing cells among CD8+ T cells. (**M-N**) Quantification of CD8+ T cell immune response in the tumor of mice + MLN by flow cytometry. The proportion of (**M**) IFN-γ and (**N**) granzyme B (GzmB) producing cells among CD8+ T cells. For all experiments, points represent results from individual animals. Bars represent the mean + SEM. Statistical analysis by Mann-Whitney test. *P<0.05, **P<0.01, ***P<0.0.001, ****P<0.0001.

As surgical resection of MLN attenuated response to ICT, we further investigated the impact of MLN removal on anti-cancer immune responses (fig. 3F). We first observed that immune cell (CD45+ cells) infiltrate in TDLNs was significantly lower in mice lacking MLN, even after ICT (fig S9A). Accordingly, TDLN weight was decreased as well (fig. S9B). MLN-resection resulted in a significant decrease in CD11c+ dendritic cells (fig. S10) as well as activated effector CD4+ and CD8+ T cell subsets (CD69+ and CD62L-) in TDLNs (fig. 3G-J, fig. S11). Indeed, the proportion of TDLN CD8+ T cells producing IFN-γ (fig. 3K) and Granzyme B (fig. 3L) was significantly decreased in mice lacking MLN (fig S12). Further, MLN removal resulted in significantly lower leukocyte cell infiltration into the tumor (fig. S13) and a concomitant reduction in the proportion of tumor CD8+ T cells producing IFN-γ (fig. 3M) and Granzyme B (fig. 3N) as well (fig S14). These data suggest that MLNs likely play a critical role in mediating the gut microbiota-dependent extra-intestinal anti-tumor immune effects observed with ICT, perhaps by functioning as gateway by which gut microbiota-induced immune signals are delivered to more distal body niches.

### Dendritic cells facilitate gut microbiota translocation into secondary lymphoid organs

We then investigated potential mechanisms by which gut microbiota translocation into secondary lymphoid organs was facilitated in the context of ICT. While combination anti-PD-1 and anti-CTLA-4 therapy has been shown to be clinically effective against various tumor types, the high incidence of developing immune-related adverse events, such as colitis, limits widespread use, particularly in older patients with multiple medical comorbidities (*18*). Thus, we explored the possibility that ICT-induced gut inflammation could lead to a disruption of intestinal epithelial barrier function thus promoting gut microbiota translocation (*32*). We orally administered fluorescein isothiocyanate (FITC)–dextran, as a means to measure gut permeability, to tumor-bearing mice treated with or without ICT. ICT did not increase the serum FITC-dextran levels during the course of ICT (fig. S15B). Further, ICT did not alter the mRNA expression of ZO-1, an intestinal epithelial tight junction protein, in the colons of mice (fig. S15C). These data suggest that increased gut permeability may not be a mechanism of ICT-induced gut microbiota translocation in this preclinical model.

Next, leveraging the knowledge that DCs can retain live commensal gut microbiota for several days and carry or traffic microbiota to MLNs (*33*), we hypothesized that the ICT-induced gut microbiota translocation could be a DC-dependent phenomenon. To test this hypothesis, we utilized CD11c-dtr transgenic mice, in which the CD11c promoter (*ltgaz*) directs the expression of a diptheria toxin receptor (dtr), and administration of diptheria toxin allows for depletion of DC populations (*34*). Indeed, a single dose of intraperitoneal diphtheria toxin (DT) injection efficiently depleted CD11c+ DC populations from MLNs (fig. 4A and B). We then measured the total bacterial load in MLNs of wild-type and CD11c-dtr mice (as determined by quantifying genomic copies of 16s ribosomal RNA gene within the total genomic DNA extracted from MLNs of mice treated with or without ICT and DT). We observed that ICT induces bacterial translocation into MLNs in wild-type mice but not in CD11c-dtr mice treated with DT (fig. 4C). This result suggests a potential key role for DCs in ICT-induced bacterial translocation into secondary lymphoid organs.

**Figure 4.**
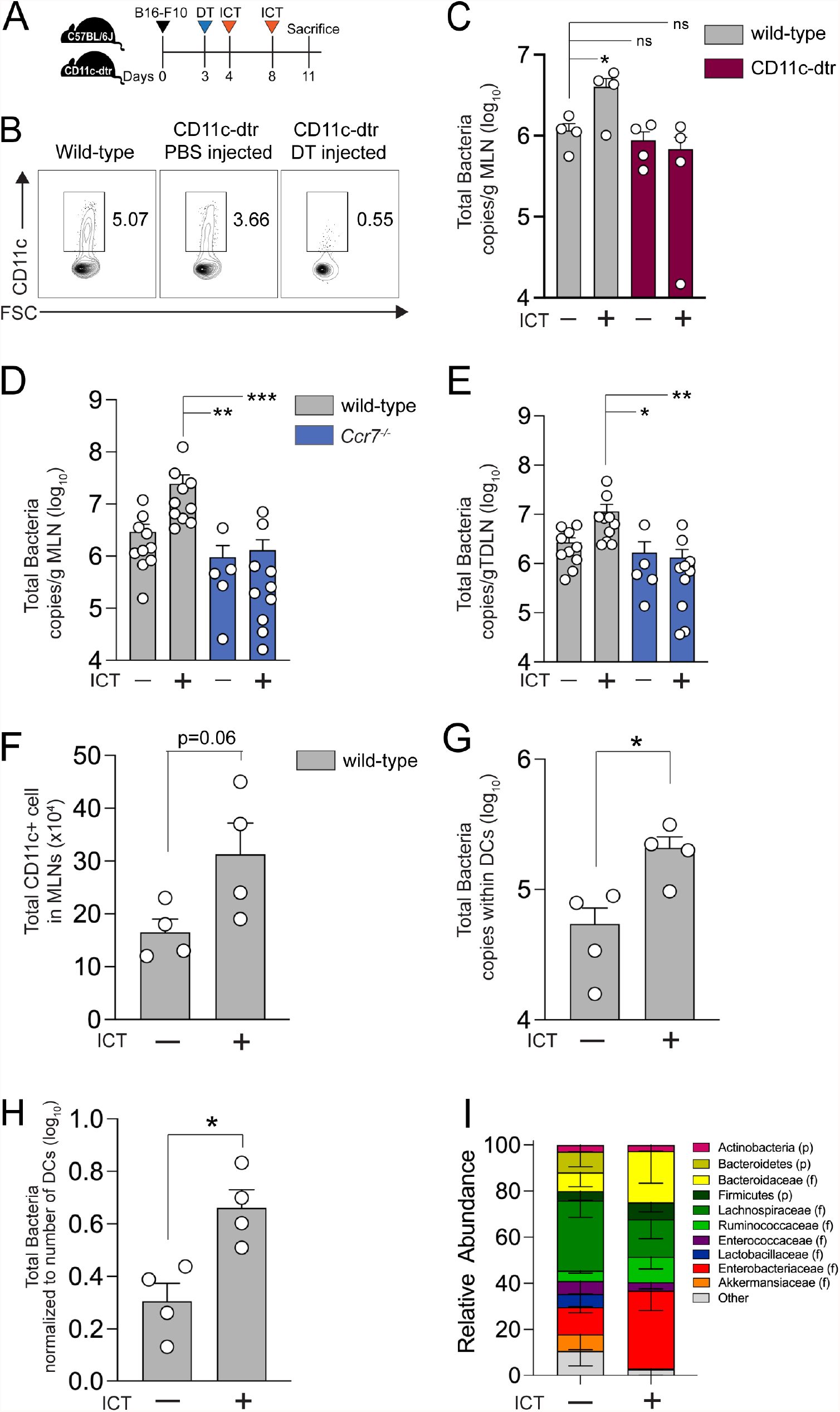
DCs facilitate ICT-induced gut microbiota translocation into secondary lymphoid organs. (**A**) Schematic overview of the protocol used to assess the impact of dendritic cell (DC) depletion on ICT-induced microbial translocation into MLN. CD11c-dtr mice (female, 6-8 wks, Jackson) were injected with 100 ng diphtheria toxin (DT) intraperitoneally on day 3 post tumor implantation to deplete CD11c+ DCs. Wild-type C57BL/6 and DT-treated CD11c-dtr mice were injected with 200μg anti-PD-1 and 200μg anti-CTLA-4 mAb intraperitoneally on days 4 and 8 post tumor implantation. (**B**) Representative flow cytometry plot of CD11c+ subsets. (**C**) Bacterial load of MLN in wild-type or CD11c-dtr DC-depleted mice + ICT, as determined by bacterial group (Eubacteria, all bacteria) quantitative-PCR of MLN gDNA collected from mice as detailed in fig. 5A. n=4 per group. Bacterial load in (**D**) MLN and (**E**) TDLN recovered from wild-type (C57BL/6J) and *Ccr7*^-/-^ mice + ICT. n=10 per group. (**F-H**) Bacterial load in DCs. CD11c+ DCs were isolated from MLN of C57BL/6 mice (female, 6-8 wks, Jackson) bearing B16-F10 melanoma tumors + ICT. gDNA was isolated from DCs (from 5 mice pooled into one sample). Bacteria load was determined by bacterial group (Eubacteria, all bacteria) quantitative-PCR of DC gDNA. (**F**) Number of CD11c+ DCs isolated from MLN of wild-type mice + ICT. (**G**) Quantification of bacterial load within DCs isolated from MLN of mice + ICT. (**H**) Quantification of bacterial load within DCs normalized to the total number of DCs. (**I**) Relative abundance of microbiota in dendritic cells, as determined by 16S rRNA sequencing. For (**D, E**), points represent results from individual animals. For (**F-H**), each point represents a single biological replicate with DCs pooled from 5 mice. Bars represent the mean + SEM. Statistical analysis by Mann-Whitney test. *P<0.05, **P<0.01, ***P<0.0.001, ****P<0.0001.

CCR7 has been shown to be important for the entry of lymphocytes and DCs into secondary lymphoid organs. Hence, we assessed the role of CCR7 in ICT-induced gut microbiota translocation into secondary lymphoid organs (*35, 36*). Wild-type C57BL/6J and *Ccr7*^-/-^ mice were implanted with melanoma tumors and treated with ICT. Consistent with our previous results, the increase in ICT-induced bacterial translocation into the MLNs and TDLNs was not observed in *Ccr7*^-/-^ mice (fig. 4D and E). Prior studies have reported that augmenting inflammation in the gastrointestinal tract (e.g. *Rnf5*^-/-^ mice (*37*) or *Salmonella* infection (*36*)) increases DC mobilization into MLNs and TDLNs in a CCR7 dependent manner. Indeed, mice treated with ICT had increased expression of the pro-inflammatory genes *Tnfa* and *Il1b* in the small intestine and colon (fig. S16). Thus, ICT-induced gut inflammation may play a role in promoting DC mobilization, and thus trafficking of gut microbiota, into secondary lymphoid organs in a CCR7 dependent fashion.

Finally, to further confirm the ability of gut microbiota to reside within DCs (*33*), we isolated CD11c+ DCs from the MLN of wild-type mice treated with or without ICT and measured total bacterial load. Not only was the total number of CD11c+ cells higher in MLNs of mice treated with ICT (fig. 4F), but the total bacterial load was significantly higher within the DCs isolated from MLNs in mice treated with ICT as well (fig. 4G). Even when normalizing to the number of DCs recovered, the total bacterial load per DC was higher in mice treated with ICT (fig. 4H). To determine the identity of the bacteria trafficked by DCs, we performed 16S rRNA sequencing on gDNA extracted from MLN DCs in melanoma bearing mice treated with or without ICT. The identified taxa in DCs (fig. 4I) was commensurate with the tissue and tumor microbiomes shown previously (fig. 1D and E). Interestingly, Enterobacteriaceae were significantly enriched in MLN DCs in mice treated with ICT (fig. S17), findings which may partially explain our prior observation that Enterobacteriaceae (*Shigella*) was significantly enriched in tumor and secondary lymphoid organ tissues of mice treated with ICT despite low relative abundance in the gut (fig 1F, S3). Collectively, these results suggest DCs play an integral role in ICT induced gut microbiota translocation mediated anti-tumor immune effects.

### Antibiotic treatment results in decreased gut microbiota translocation into MLN, decreased polyfunctional CD8+ T cell effector responses, and diminished ICT efficacy

Antibiotic exposure has been associated with inferior clinical outcomes in cancer patients receiving ICT (*6, 12, 38-40*). Germ-free mice and mice pre-treated with antibiotics are less responsive to ICT (*13, 14*). Yet, the mechanisms by which antibiotic treatment attenuates effectiveness of ICT are unclear. We posited that antibiotic treatment would reduce overall levels of gut microbiota, thus reducing gut microbiota translocation into secondary lymphoid organs and tumor, thereby mitigating gut microbiota-induced immune activation and ultimately attenuating ICT efficacy. We treated mice with or without antibiotics one week before tumor implantation and then proceeded with ICT (fig. 5A). Consistent with prior reports (*12-14*), antibiotic treatment induced an ICT hyporesponsive state in our preclinical cancer model (fig. 5A). Further, antibiotic exposure significantly reduced gut microbiota translocation into MLNs (fig. 5B). To further investigate the immunological impact of antibiotic-induced gut microbiota depletion, we isolated CD8+ T cells from MLNs and TDLNs of control or antibiotic treated mice and performed single-cell multiplex cytokine profiling (Isoplexis IsoSpark; 28-plex mouse adaptive immune IsoCode chip panel). Overall, the MLN and TDLN CD8+ T cell cytokine secretion profile was markedly distinct between the antibiotic treated mice versus untreated controls (fig. 5C and D). Specifically, the number of CD8+ effector T cells (defined as T cells secreting IFN-γ, granzyme-B, MIP1a and/or TNF-α) was significantly reduced in antibiotic-treated mice (fig. 5E, S18, 5F, S19). Interestingly, T-cell polyfunctionality, the ability of T-cells to secrete two or more cytokines per cell, has been associated with positive clinical response to cancer immunotherapies (including ICT and CAR-T therapy) in mice and humans (*41-44*). Indeed, MLN and TDLN CD8+ T cells recovered from mice without antibiotic exposure and treated with ICT exhibited higher T-cell polyfunctionality (fig. S20). CD8+ T cells secreting both granzyme-B and interferon-γ were more frequently observed in untreated mice (fig. S18 and S19). Accordingly, the number of MLN and TDLN CD8+ T cells secreting interferon-γ (IFN-γ) or granzyme-B (GZMB) was significantly lower in antibiotic-treated mice compared to untreated controls (fig. 5G and H). These results suggest that antibiotic-induced depletion of endogenous gut microbiota and a subsequent decrease in gut microbiota translocation results in reduced ICT efficacy via altering CD8+ T cell effector responses in secondary lymphoid organs.

**Figure 5.**
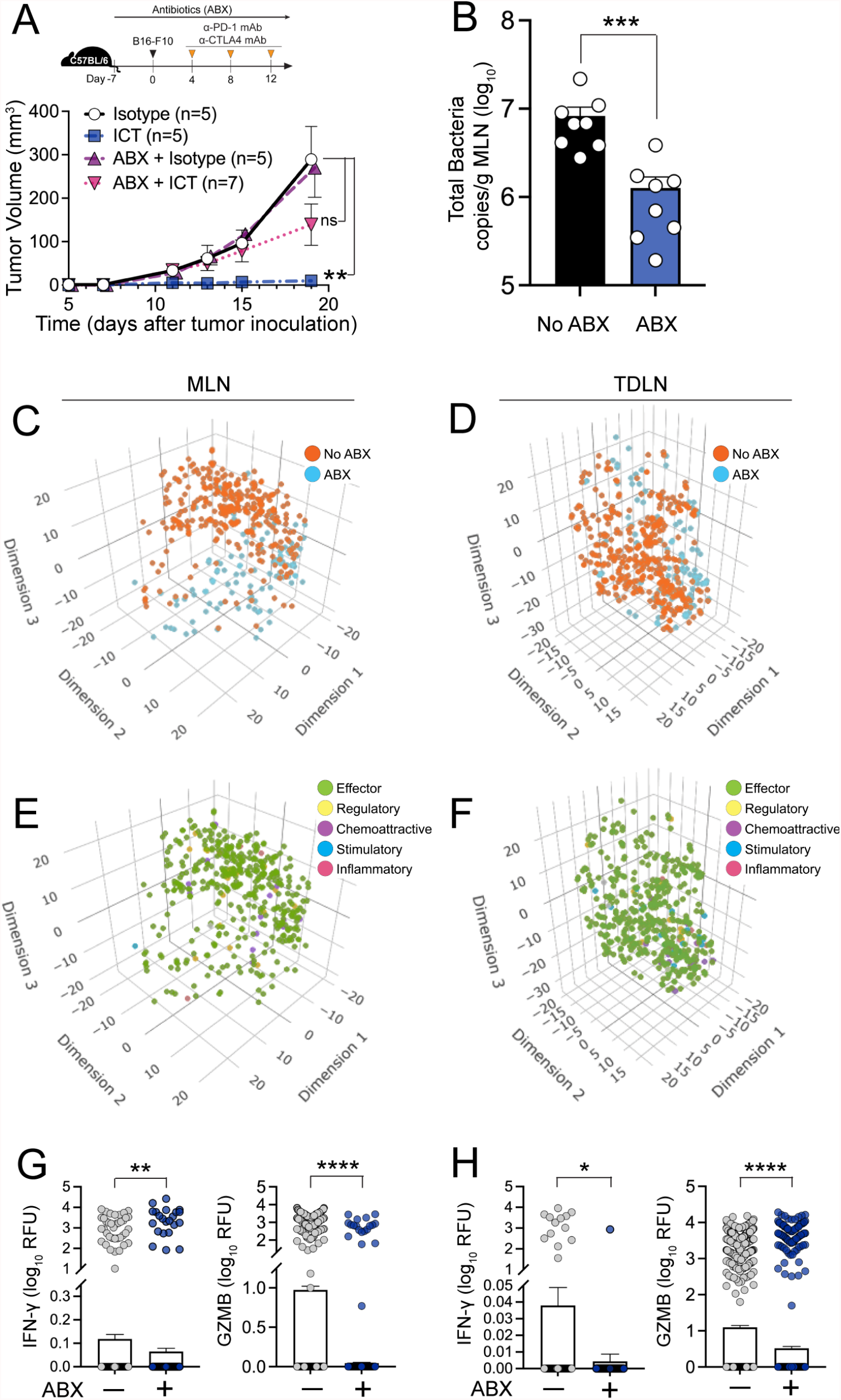
Antibiotic treatment results in decreased gut microbiota translocation into MLN, decreased polyfunctional CD8+ T cell effector responses, and diminished ICT efficacy. Schematic overview of the protocol used to assess the impact of antibiotic-induced gut microbiota depletion on ICT efficacy, gut microbiota translocation and effector CD8+ T cell response. C57BL/6 mice (female, 6-8 wks, Jackson) were treated + antibiotics (ABX, 2 mg/ml streptomycin and 1500 U/ml penicillin G in drinking water) for 7d before B16-F10 tumor inoculation. Mice were treated with 200μg anti-PD-1 and 200μg anti-CTLA-4 mAb intraperitoneally on days 4, 8, and 12 after tumor implantation (A) Tumor volume of mice + ABX and ICT. n=5-7 mice per group. (B) Bacterial load of MLN in mice + ABX and ICT, as determined by bacterial group (Eubacteria, all bacteria) quantitative-PCR of MLN gDNA collected from mice as detailed in fig. 4A. n=8 per group. Three-dimensional t-distributed stochastic neighbor embedding (t-SNE) plot of secretory cytokine profiles of CD8+ T-cells isolated from (**C**) MLN (n=3 per group) and (**D**) TDLN (n=10 per group) of mice + ABX + ICT, as determined by using Isoplexis IsoSpark; 28-plex mouse adaptive immune panel. Only CD8+ T cells secreting at least one cytokine are represented. tSNE plot displaying immune functional categories of CD8+ T cells from (**E**) MLN and (**F**) TDLN, defined as Effector: Granzyme B, IFN-γ, MIP-1α, TNF-α Stimulatory: GM-CSF, IL-12p70, IL-15, IL-18, IL-2, IL-21, IL-5, IL-7 Chemoattractive: BCA-1, CCL-11, IP-10, RANTES, CXCL1, CXCL13 Regulatory: FAS, IL-10, IL-13, IL-27, IL-4, sCD137 Inflammatory: IL-17A, IL-1β, IL-6, MCP-1 Absolute quantification of IFN-γ and Granzyme B secreting CD8+ T cells isolated from (**G**) MLN and (**H**) TDLN of mice + ABX + ICT. For (**B**), each point represents individual animals. For (**C, D, E, F**), each point represents individual CD8+ T cells isolated from MLN and TDLN of mice + ABX + ICT. Bars represent the mean + SEM. Statistical analysis by Mann-Whitney test. *P<0.05, **P<0.01, ***P<0.0.001, ****P<0.0001.

## Discussion

We have identified gut microbiota translocation into secondary lymphoid organs as a general mechanism by which resident bacteria in the gut can shape and dictate extra-intestinal anti-tumor immune responses in the setting of ICT. ICT enhances the ability of DCs to facilitate gut microbiota translocation into secondary lymphoid organs. And specific gut microbiota taxa have a greater predilection or capacity to translocate, with a notable differential ability of some microbiota taxa to induce anti-cancer immune responses.

Numerous mechanisms are being posited by which gut microbiota can enhance anti-tumor responses in the setting of ICT: gut microbiota expressing peptide sequences homologous to tumor antigens/neoantigens (molecular mimicry) (*21*); gut microbiota-derived metabolites (SCFA (*3*), inosine(*9*)) ; or direct activation of innate/adaptive immune cells to drive anti-tumor responses (*20, 45*). All of the aforementioned mechanisms require gut microbiota or gut microbiota-derived product engagement with host innate and/or adaptive immune effectors. The emerging concept of the tumor microbiome (*46, 47*) provides one possible explanation: resident microbiota or microbiota-derived metabolites within the tumor inducing anti-tumor effects. Of note, the relative abundance of Bacteroides (10-15%) and Firmicutes (40-70%) in our murine melanoma tumors are comparable to those observed in tumor microbiomes in human melanoma patients, 15% and 40% respectively (*46*). In addition, the highly abundant translocator taxa that we identified in our study, (Enterococcaceae, Lactobacillaceae, and Enterobacteriaceae) were also detected in melanoma tumors in humans (*46*). More intriguingly, these taxa identified in both murine and human specimens originate from the gut, as opposed to from adjacent tissues such as the skin. But an unanswered question is how do gut microbiota end up in such distal sites as the skin or lung?

One potential explanation is gut microbiota translocation. The ability of gut microbiota to translocate into secondary lymphoid organs, particularly MLN, has been well-described in infectious (*48-51*) and inflammatory diseases (*52-54*). Further, a seminal prior study highlighted the importance of gut microbiota translocation into secondary lymphoid organs for optimal cyclophosphamide immune-mediated anti-cancer responses, identifying a novel mechanism by which this commonly used alkylating chemotherapeutic agent induces host immune anti-cancer effects (*22, 55*). Our data suggest that an analogous process may be one mechanism by which ICT promote gut microbiota translocation into secondary lymphoid organs: ICT-induced gut inflammation creating an environment in which gut microbiota are more readily able to translocate into secondary lymphoid organs to engage innate and adaptive immune effectors.

Interestingly, while bacteremia, and the development of sepsis, is generally associated with increased morbidity and mortality, it has been associated with improved cancer outcomes in both preclinical models (*56*) and in patients (*57, 58*). And localized infections, proximal to primary tumors, have also been associated with improved clinical outcomes in animals (dogs with osteosarcoma (*59*)) and humans (head and neck cancer (*60*) and osteosarcoma (*61*)). Yet the development of bacteremia has not been associated with ICT treatment. And our data did not identify any appreciable deficit in gut barrier integrity with ICT. Nonetheless, in addition to gut microbiota translocation into secondary lymphoid organs, a low-grade and subclinical gut microbiota translocation into the systemic circulation could be an additional potential mechanism by which tumor microbiomes develop.

While the role of DCs in anti-cancer immunity is well-established, our data provide an additional layer of insight in that gut microbiota-dependent anti-cancer immune effects may also be dependent on DCs capacity to carry or traffic gut microbiota. Previous studies have reported that gut inflammation increases DC mobilization into MLNs and TDLNs in a CCR7 dependent manner (*36, 37*). Our data corroborate these findings and also add that the degree of gut microbiota translocation (bacterial burden) into secondary lymphoid organs is DC dependent. A recent study reported the frequent intracellular localization of microbes in cancer and immune cells (*46*), thus further supporting the role of DCs in bacterial translocation

In summary, by illuminating a general mechanism by which gut microbiota shape or influence extraintestinal anti-cancer immune responses, our results could provide insight as to why different gut microbiota taxa or mechanisms are being espoused as critical for ICT efficacy. Ultimately, multiple gut microbiota taxa and/or mechanisms have the potential to drive host anti-cancer immune responses, but one common prerequisite is that these microbes or microbe-derived metabolites must engage with key components of the innate and adaptive immune systems. As such, ICT and DC-mediated gut microbiota translocation into secondary lymphoid organs could serve as a fundamental mechanism by which these processes transpire. In the future, these principles could also apply to understanding if and how gut microbiota modulate the efficacy of other cancer immunotherapies, as these general insights are neither immunotherapy or tumor specific.

## Author contribution

Y.C., L.V.H, C.P., and A.Y.K, designed the research; Y.C., Y.G., L.A.C., and N.P performed experiments; Y.C., J.K., X.Z., and A.Y.K. analyzed data; Y.C, L.V.H, C.P., and A.Y.K edited the manuscript; Y.C. and A.Y.K wrote the paper; and all authors revised the manuscript and approved its final version.

