## Supplementary material for "Immune checkpoint blockade induces gut microbiota translocation that augments extraintestinal anti-tumor immunity": Choi_et_al_Supplementary_Materials.pdf

**This PDF file includes:**

Materials and Methods
Figures S1-S20
Table S1-S3

### Materials and Methods

**Mice**. C57BL/6J (Stock No: 000664), B6.FVB-1700016L21RikTg (Itgax-DTR/EGFP)57Lan/J) (CD11c-DTR, Stock No: 004509) (1), C57BL/6-Tg(TcraTcrb)1100Mjb/J (OT-1, Stock No: 003831) (2), and B6.129P2(C)-Ccr7tm1Rfor/J (CCR7-/-, Stock No: 006621) (3) mice were obtained from Jackson Laboratories and bred and maintained in the barrier facility at the University of Texas Southwestern Medical Center. All animals were kept on a 12-hour light-dark cycle and were fed standard mouse chow (Teklad 2916, irradiated). Sex-matched, 6-8 week old mice were used for all experiments and co-housed littermates were used as controls. Experiments were performed using protocols approved by the Institutional Animal Care and Use Committees of the UT Southwestern Medical Center.

**Preclinical model of melanoma and immune checkpoint therapy.** B16-F10 cells (ATCC CRL-6475; RRID:CVC_0159) were grown at 37°C under 5% $CO_2$ in DMEM medium supplemented with 10% heat-inactivated FBS (Sigma), 100 units/ml penicillin, 100μg/ml streptomycin sulfate, 2mM L-glutamine, 1mM sodium pyruvate (Thermo Fisher). C57BL/6J mice were fed sterile or antibiotic-supplemented (2 mg/ml streptomycin and 1500 U/ml penicillin G, Sigma). $1 \times 10^5$ B16-F10 cells were implanted subcutaneously into the right flank of mice. Four days after the tumor inoculation, mice were injected with 200μg anti-PD-1 antibody (RMP1-14, CD270, BioXcell) and anti-CTLA-4 antibody (9D9, CD152, BioXcell) or isotype control (Rat IgG2a, κ or Mouse IgG2b, respectively) intraperitoneally; an additional two treatments were given with 4d intervals. Tumor volumes were calculated using measurements from a digital caliper and the following formula: $\pi/6$ x length x width x height (4). Loss of survival was defined as death (with moribund mice being euthanized) or when tumor diameter > 2 cm in any dimension.

### Gut and Tissue Microbiome Profiling

***Cultured Microbiota Enumeration.*** Mice were euthanized and tissue samples (mesenteric lymph nodes (MLN), tumor draining lymph node (TDLN), spleen, and tumor) were resected and placed into a pre-weighed screw-cap microfuge tube (2ml, Fisher Scientific) with 500ul sterile reduced PBS and one sterile 5mm borosilicate glass beads (Sigma). Samples were kept in a 2.5L portable anaerobic chamber (Mitsubishi Gas Chemical) with anaerobic gas packs (Mitsubishi Gas Chemical) during sample collection and transportation. Tissue samples were homogenized using TissueLyser II (Qiagen) at 30 Hz for 3 minutes. Tissue homogenates were serially diluted in reduced PBS and plated on BHI/Blood, YCFA, and CME0151 agar and incubated for 24-72 hours at 37°C under anaerobic conditions (Coy anaerobic chamber). Colony-forming units (CFUs) were counted. Representative colonies based on morphology (~10-20 for each morphology type) were picked and cultured in YCFA medium (Anaerobe Systems) for 24-48 hours at 37°C under anaerobic conditions. Liquid cultures of representative colonies were centrifuged at 4,000g for 10 min at 4°C. Bacterial pellets were suspended in 710 ul of extraction buffer (200 mM NaCl, 200 mM Tris, 20 mM EDTA, 6% SDS) and 0.5 ml of phenol-chloroform–isoamyl alcohol, pH 7.9 (Ambion). Cells were lysed by bead-beating (0.1 mm zirconia/silica beads for bacterial gDNA (BioSpec) and subjected to additional phenol-chloroform extractions. Crude DNA extracts were treated with RNaseA (Qiagen) and column-purified (PCR Purification Kit, Qiagen). DNA concentrations were quantified by a fluorescence-based assay (Quant-iT PicoGreen dsDNA, Life Technologies). V1-V9 region of the 16S rRNA gene was then amplified from each cultured sample using the following primers. Forward primer 27FMod, 5'-AGRGTTTGATYMTGGCTCAG-3' and reverse primer 1492R, 5'-GGYTACCTTGTTACGACTT-3'. Accuprime Pfx Supermix was used for PCR reaction (Invitrogen). Reaction conditions were 5 min at 95°C, followed by 35 cycles of 15 s at 95°C, 30 s at 55°C, 90 s at 68°C, and a hold at 4°C on an Eppendorf Mastercycler. Amplicons were then sent for Sanger sequencing. Sequences were entered into the NCBI standard nucleotide Basic Local Alignment Search (BLAST) tool utilizing the rRNA/ITS databases. Bacterial species identification was ascertained from BLASTN results with the highest Total Score, with percent identity score >95% and E value <0.01.

***16S rRNA sequencing and analysis.*** gDNA was extracted from fecal and tissue (MLN, spleen, TDLN, and tumor) samples using the MagAttract Power Microbiome DNA/RNA KF kit (Qiagen) and Kingfisher Flex (Thermo Fisher Scientific). 16S rRNA genes (variable region 4, V4) were amplified from each sample in 96-well plates using a composite forward primer and a reverse primer containing a unique 8-base barcode that was used to tag PCR products from respective samples (*5*). We used the reverse primer 926R, 5'-*CAAGCAGAAGACGGCATACGAGAT*-NNNNNNNN-**AGTCAGTCAG-CC-GGACTACHVGGGTWTCTAAT**-3": the italicized sequence is the reverse MiSeq primer i7; NNNNNNNN designates the unique 8-base barcode used to tag each PCR product; the bold sequence is the broad-range 16S bacterial primer containing the pad-link-16SR. The forward primer used was 515F, 5'-*AATGATACGGCGACCACCGAGA TCTACAC*-NNNNNNNN-**TATGGTAATT-GT-GTGCCAGCMGCCGCGGTAA**-3': the italicized sequence is MiSeq Primer i5; the NNNNNNNN designates the unique 8-base barcode used to tag each PCR product; and the bold sequence is the broad range 16S bacterial primer containing the pad-link-16SF. PCR reactions consisted of 17ul Accuprime Pfx Supermix, 1000 nM of each primer, and 20ng of template. Reaction conditions were 2 min at 95°C, followed by 30 cycles of 20 s at 95°C, 15 s at 55°C, 5 min at 72°C, then 10 min at 72°C, and a hold at 4°C on an Eppendorf Mastercycler. For tissue and tumor microbiome profiling, two rounds of PCR amplification were performed. Products were verified on a 1% agarose gel, and all samples were cleaned and normalized using the AmPure Normalization plate protocol using the KingFisher Flex platform. Each plate was then pooled into a single tube, and the PCR product size and library quality of each individual pooled plate was checked using Agilent Technologies D1000 ScreenTape electrophoresis. Additionally, KAPA Biosystems PCR Library Quantification kit was used to quantify each pooled plate. Illumina spike-in (PhiX) was included at 4 pM at 10%, and the pooled sample library was included at 4pM at 90% yielding a final library concentration of 3.6 pM and PhiX concentration of 0.4 pM. Pooled samples were then sequenced with an Illumina MiSeq (PE-250). Raw sequences generated from Illumina MiSeq for 16S rRNA gene PCR amplicons were quality filtered. Sequences shorter than 200 nucleotides or longer than 1000 nucleotides were removed. Sequences containing ambiguous bases, primer mismatches, homopolymer runs in excess of 6 bases and uncorrectable barcodes were also removed. Sequences that passed the quality filtration were denoised and analyzed using the open source software package Quantitative Insights Into Microbial Ecology (QIIME2). 16S rRNA gene sequences were then classified taxonomically using the classifier Silva database. Differential taxonomic abundance between different phenotypic groups was analyzed by linear discriminate analysis (LDA) coupled with effect size measurements (LEfSe). Only taxa noted to have >2 log fold increase in LDA score and $p < 0.05$ (Kruskal-Wallis test) were identified as significantly enriched or depleted.

**Quantitative PCR for tissue microbiome analysis.** Bacterial load in tissue samples was quantified by qPCR analysis (SsoAdvanced SYBR Green Supermix, Bio-Rad) of microbial gDNA using the universal 16S rRNA gene. Bacterial abundance was determined using standard curves constructed with reference to cloned DNA corresponding to a short segment of the 16s rRNA genes that were amplified using conserved specific primers, forward, 5'- ACTCCTACGGGA-GGCAGCAGT-3', and reverse, 5'- ATTACCGCGGCTGCTGGC-3'. 25ng of microbial gDNA was used as an input template. Reaction conditions were 30 sec at 95°C, followed by 40 cycles of 1 sec at 95°C and 10 sec at 63°C on a BioRad CFX96 Real-Time System. Total 16s copy number per sample was determined by standard curve and normalized to the tissue weight. Note that qPCR measures gene copies per gram of tissue, not actual bacterial numbers or colony-forming units. For the quantification of bacterial load in isolated dendritic cells, CD11c+ DCs isolated from MLNs of five mice were pooled together. Pooled CD11c+ DCs were then suspended in gDNA extraction buffer and proceeded with the gDNA extraction protocol described above.

**Surgical removal of secondary lymphoid organs.** $1 \times 10^5$ B16-F10 cells were implanted subcutaneously into the right flank of C57BL/6J mice. On day 7 post tumor implantation, mice with tumor volumes of 100 $mm^3 \pm 20$ $mm^3$ were then randomized to one of the following groups: 1) mesenteric lymph nodes (MLN) resection; 2) spleen resection; 3) inguinal lymph node (TDLN), defined as the right inguinal lymph nodes as all tumors were implanted in the right flank; and 4) sham, longitudinal abdominal incision only. Mice were anesthetized with vaporized isoflurane. Abdominal fur was removed using an electric razor followed by Veet hair removal cream (Veet). The abdomen was then cleaned with betadine x 3 and 70% ethanol. Mice were then covered with sterile surgical drapes with a Glad Press N Seal (Glad) over the mid-abdominal surgical area site. A longitudinal abdominal incision was then performed. Intestines were gently removed from the peritoneal cavity and placed on a sterile gauze pads (moistened with sterile normal saline solution). Target lymphoid organs (mesenteric lymph node, inguinal lymph node or spleen) were carefully resected with surgical scissors and/or a cauterizer (Medline). Large blood vessels were cauterized and ligated. The abdominal wall was closed using absorbable sutures (5/0 PGA, Covetrus) in individual stitches and then the skin is closed using nonabsorbable sutures (5/0 Monofilament nylon, Covetrus) and tissue adhesive (veterinary surgical adhesive, Covetrus). The incision was then coated with triple antibiotic ointment (Bacitracin zinc, neomycin sulfate, and polymyxin b sulfate ointment, Taro pharmaceutical). Post-operative analgesic carprofen (5 mg/kg) was administered intraperitoneally immediately after surgery. Anesthesia was discontinued. Mice were initially placed on a surgical bed for recovery, but once moving transitioned to a heated cage. Mice were monitored closely and evaluated for pain and lethargy. Any moribund mouse (e.g. signs of lethargy, cool to touch, etc.) was immediately euthanized. At a 4-hour post-surgery check, buprenorphine (0.05 mg/kg SQ) was administered as needed for pain. For the first 72 hours, mice were monitored twice daily for pain management and infection monitoring. Mice found dead or euthanized for being moribund within the first 72 hours after surgery were not included in the final analysis, as mortality was attributed to either post-surgical bleeding or infection.

**Single cell suspension preparation from secondary lymphoid organs.** Dissected mesenteric lymph nodes and inguinal lymph nodes (tumor-draining lymph nodes) were kept in ice-cold sterile PBS until processing. Tissue samples were placed on a 70 µm sterile cell strainer (Fisher) and mashed with the plunger of a 10 ml syringe (BD) over a sterile petri dish (Falcon). An additional 5 ml of ice cold-was added to the cell strainer. Cell suspension was then collected from the petri dish and transferred into a 15ml conical tube. Cell strainer and plunger were washed with additional 10 ml of ice-cold PBS, and the flow-through was collected and pooled to the cell suspension. The cell suspension was centrifuged at 300 G, 4°C for 10 min. The supernatant was carefully aspirated, and the cell pellet was suspended in ice-cold PBS for further downstream use.

**Tumor infiltrating lymphocytes isolation.** Single cells from dissected tumor tissue were isolated using gentleMACS™ Dissociators and mouse tumor dissociation kit, according to the manufacturer's protocol (Miltenyi Biotec). Briefly, the tumor was cut into 2–4 mm pieces and mechanically dissociated with gentleMACS™ Dissociators. Dissociated tumor tissue was then enzymatically digested using tumor dissociation kit enzyme D, R and A at 37°C for 40 min (Miltenyi Biotec). Tumor tissue was further dissociated using gentleMACS™ Dissociators and passed through a 70 µm sterile cell strainer (Fisher). The single-cell suspension was then transferred into a 15ml conical tube and washed with 10 ml of RPMI medium. The cell suspension was centrifuged at 300 G, 4°C for 10 min. The supernatant was carefully aspirated, and the cell pellet was suspended in RBC lysis buffer (Invitrogen) and incubated at room temperature (RT) for 2 min to remove erythrocytes. The cells were washed (10ml of PBS) and centrifuged at 300 G, 4°C for 10 min. Cell pellets were suspended in 40% Percoll solution and carefully transferred to 15ml conical tubes containing 80% Percoll solution. Suspensions were centrifuged at 300 G at RT for 20 min (ascending rate: 5; descending rate: 0). Cells at the interface between 40% and 80% Percoll solutions were carefully collected and washed once with PBS.

**Flow cytometry analysis.** Single-cell suspensions of cells were transferred to U-bottom 96 well plate (Corning) and centrifuged at 300 G, 4°C for 10 min. For surface staining, cells were incubated with zombie-yellow live/dead dye (Biolegend) diluted in PBS (1:500) at RT for 15 minutes. After washing with PBS, cells were incubated with Fcγ receptor blocking antibody (clone 2.4G2, BD Biosciences), diluted in FACS buffer, PBS supplemented with 2% heat-inactivated fetal bovine serum (FBS) and 2mM EDTA, for 10 min at 4°C in the dark followed by surface staining for 30 min at 4°C in the dark. Cells were then washed with FACS buffer twice and suspended in FACS buffer for flow cytometry. For intracellular staining, cells were incubated in 200μl RPMI supplemented with 10% FBS, 50 μM 2-mercaptoethanol, 50ng/ml Phorbol 12-Myristate 13-Acetate (PMA), 750ng/ml Ionomycin and 10μg/ml Brefeldin-A (all Sigma) at 37°C with 5% CO2 for 4 hours. After the incubation, cells were washed with PBS. Cells were then incubated with zombie-yellow live/dead dye (Biolegend) diluted in PBS (1:500) at RT for 15 minutes. After washing with PBS, cells were incubated with Fcγ receptor blocking antibody (clone 2.4G2, BD Biosciences), diluted in FACS buffer for 10 min at 4°C in the dark followed by surface staining for 30 min at 4°C in the dark. Cells were then fixed and permeabilized using the eBioscience™ Foxp3 Fixation/Permeabilization kit (eBioscience) according to the manufacturer's protocol. Cells were then stained with antibodies targeting intracellular proteins for 30 min at 4°C. Prior to the acquisition, cells were washed with FACS buffer and flow cytometry was performed on a BD FACSLyric. Flow cytometry gating strategy for the various experiments used in this study are detailed in fig. S11, S12, and S14. Antibodies used for flow cytometry are listed in Table S2.

**Extraction of bacterial lysates.** *Enterococcus* spp. were grown in YCFA medium under anaerobic conditions (Coy anaerobic chamber) at 37 °C; and *Lactobacillus* spp. were grown in Lactobacilli MRS medium (Difco) under anaerobic conditions at 37 °C. Bacterial cultures were grown to the late log phase (OD600 = 0.8–1.2), washed with sterile PBS x 3, and suspended in sterile PBS. Bacterial suspensions were subjected to three cycles of snap-freeze-thaw in liquid nitrogen and lysed by repeated sonication. Lysates were centrifuged at 3,000 G, 4°C for 20 min to remove insoluble components, and the supernatant was filtered through a 0.22-μm membrane filter. Protein concentration was determined by Pierce BCA assay (Thermo).

**Expansion of mouse splenic DCs by B16-FLT3L implant.** For the generation of spleen-derived dendritic cells (DCs), 5 × $10^6$ B16-FLT3L cells (RRID:CVCL_IJ12) were injected subcutaneously into the right flank of C57BL/6J mice. After 10–16 d, spleens were harvested. CD11c+ DCs were isolated using CD11c microbeads and magnetic-activated cell sorting (Miltenyi). Of note, B16-FLT3L injection did not induce CD11c+ DC maturation, as reported previously (*6, 7*), as evidenced by a lack of proliferation of non-stimulated CD11c+DCs.

***ex-vivo*** **immune cell priming assay**. For the DC stimulation assays, CD11c+ DCs were stimulated with different bacterial lysates for 6 hours at 37°C under 5% CO2 and then washed with PBS. The surface expression of DC activation markers was measured by flow cytometry as described above. For T cell assays, naïve CD8+ T-cells were isolated from the spleens of OT-I (C57BL/6-Tg(TcraTcrb)1100Mjb/J) mice (female, 6-8 weeks age). Cells were maintained in complete RPMI-1640 medium (Sigma) supplemented with 10% FBS, 55µm 2-Mercaptoethanol (Sigma), 100 units/ml penicillin and 100 µg/ml streptomycin sulfate at 37°C with 5% CO2. CD11c+ DCs were pulsed with different bacterial lysates and OVA257-264 peptide for 6h followed by thorough washing, and then co-cultured with naïve OVA257-264 specific OT-I CD8+ T cells for 7 days. After co-culture, cells were washed with PBS. T cell activation and IFN-γ production were measured by flow cytometry as described above.

**Single-cell multiplex cytokine profiling of murine T cells**. Lymphocytes from MLNs and TDLNs of mice (C57BL/6J, female, 6-8 weeks old) treated ± antibiotics in the drinking water (2 mg/ml streptomycin and 1500 U/ml penicillin G) and implanted with B16-F10 tumors and treated with ICT (anti-PD1 and anti-CTLA-4 antibody treatment) were collected at one day after the last

ICT treatment, and cultured overnight in recombinant murine-IL-2 (1μg/ml, Peptrotech)-supplemented RPMI 1640 medium. CD8+ T cells were then isolated using CD8 microbeads (Miltenyi) and were stimulated with immobilized anti-mouse CD3 (Invitrogen) and soluble anti-mouse CD28 (Invitrogen) at 37 °C, 5% $CO_2$ for 48 h. Approximately 30,000 CD8+ T cells were loaded onto an IsoCode chip (IsoPlexis, New Heaven, CN) containing ~12,000 microchambers pre-patterned with a 28-plex antibody array, imaged for single cell location in microchambers and incubated at 37 °C, 5% $CO_2$ for additional 16 h. Following incubation period, ELISA detection was used to determine which combinations of proteins were being secreted by each individual cell. Secreted proteins from single cells were captured by antibody-barcoded slides; the polyfunctional profile (2+ proteins per cell) of single cells was evaluated by IsoPlexis' software.

***in vivo* Dendritic Cell Depletion.** For dendritic cell depletion experiment, CD11c-DTR mice were intraperitoneally injected with 100ng of diphtheria toxin (Sigma) as previously described (*1*). For confirming the dendritic cell depletion, spleens from the wild-type mice and CD11c-DTR mice injected with diphtheria toxin or PBS were resected 48 hours after diphtheria toxin injection and CD11c expression among the splenic lymphocytes were determined by flow cytometry.

**Gastrointestinal barrier function assays.**

***FITC-dextran permeability assay.*** Mice were fasted overnight. FITC-dextran (500 mg/kg; Sigma Aldrich; 4 kD) was administered via oral gavage. Mice were kept without food and water for 4 hours. Blood samples were obtained by terminal cardiac puncture and collected in BD Vacutainer SST tube (BD) 4 hours after FITC-dextran administration. Blood samples were centrifuged at 2,000xG at RT for 10 min. Serum was collected by taking the upper layer after centrifugation. The serum fluorescence intensity was measured at an excitation wavelength of 485 nm and an emission wavelength of 528 nm using a spectrophotometer (Synergy HT, BioTek).

**Quantitative real-time PCR (qPCR)**. Total RNA was isolated from MLN, TDLN and tumor of wild-type, CCR7-deficient, CD11c-DTR, MLN-resected mouse treated with or without ICT or antibiotics. cDNA was synthesized from purified RNA using iScript cDNA synthesis kit (Bio-Rad). qPCR was performed using SsoAdvanced Universal SYBR Green Supermix (Bio-Rad) on a CFX96 Real-Time System (Bio-Rad). Relative expression values were determined using the comparative Ct (ΔΔCt) method (*8*) and transcript abundances were normalized to 18S rRNA transcript abundance. Amplification of target genes were conducted using following primers. Mouse 18s rRNA forward: 5'-CATTCGAA- CGTCTGCCCTAT-3', mouse 18s rRNA forward reverse: 5'-CCTGCTGCCTTCCTTGGA-3' mouse ZO-1 forward: 5'-ACCCGAAACTGATGC-TGTGGATAG-3', mouse ZO-1 reverse: 5'-AAATGGCCGGGCAGAACTTGTGTA-3', mouse TNF-α forward: 5'-TCTCATGCACCACCATCAAGGACT-3', mouse TNF-α reverse: 5'-TGA-CCACTCTCCCTTTGCAGAACT-3', mouse IL-1β forward: 5'-AAGGGCTGCTTCCAAACCT-TTGAC-3', and mouse IL-1β reverse: 5'-ATACTGCCTGCCTGAAGCTCTTGT-3'.

**Human melanoma patient survival analysis with TCGA dataset.** Analysis of the intratumoral mRNA expression of toll-like receptor (TLR) and co-stimulatory receptor genes with overall survival of human skin cutaneous melanoma patients (n=458) was analyzed by utilizing the OncoLnc package and The Cancer Genome Atlas skin cutaneous melanoma (TCGA SKCM) dataset (*9, 10*). Briefly, OncoLnc incorporates TCGA SKCM dataset and calculates log-rank P values and Cox coefficients by utilizing multivariate Cox regression modeling to assess the significance. The raw output data was downloaded from http://www.oncolnc.org/.

**Data and Materials Availability**

The 16S data for this study has been deposited in the NCBI Sequence Read Archive: http://www.ncbi.nlm.nih.gov/sra/?????.

**Statistical analysis.** GraphPad Prism v.9.2 was used for statistical analysis. All data sets were tested for normality (e.g. Shapiro-Wilk). Data sets with normal distribution were analyzed with parametric tests, such as standard student t-test or one-way ANOVA with Bonferroni post-test. For non-normal distributions, non-parametric tests, such as Mann Whitney U test or Kruskal Wallis with Dunn's post-test, were applied. Two-way ANOVA with Bonferroni post-test was used for tumor growth curves. Survival was analyzed using the Mantel-Cox Log-rank test.

**Reagents and Resources.** A list of key reagents and resources used for this study are listed in Table S2.

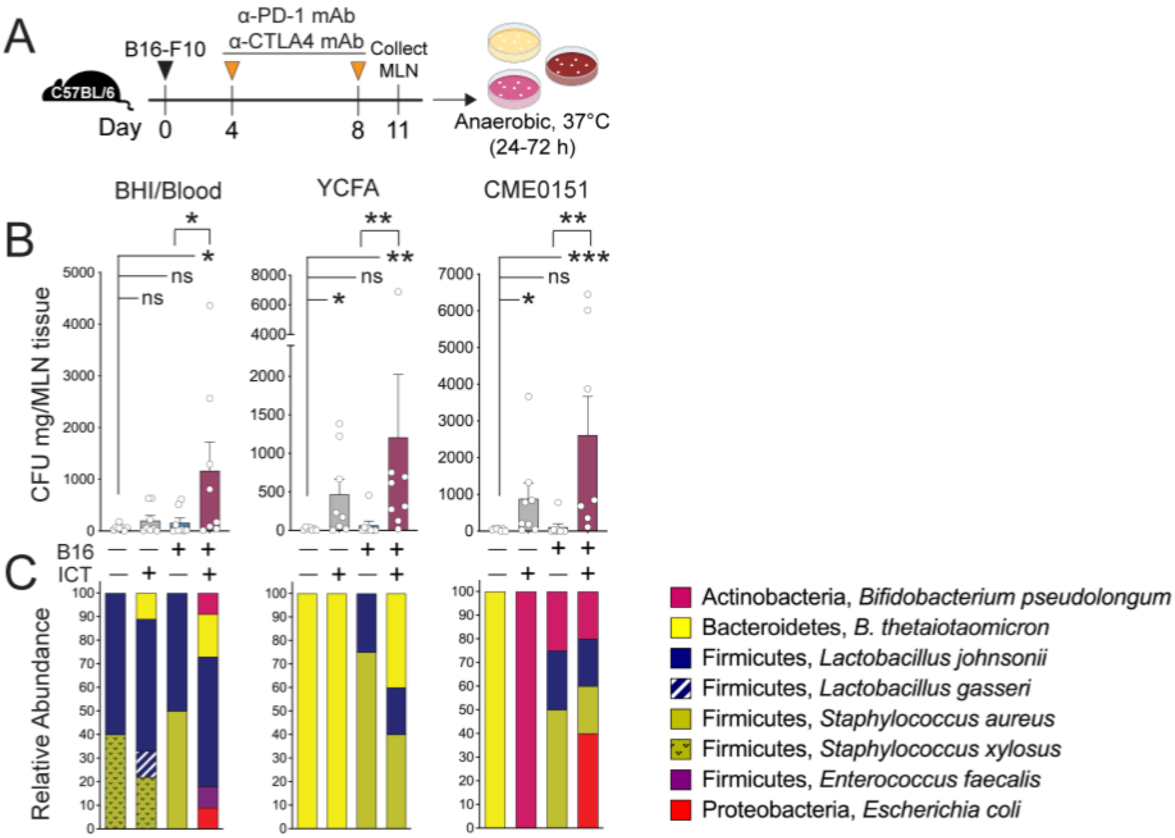


**Figure S1. Immune checkpoint inhibitor therapy (ICT) induces bacterial translocation into mesenteric lymph nodes (MLN)**

**(A)** Schematic overview of the protocol used to assess and quantify bacterial translocation into MLN in C57BL/6J mice (female, 6-8 wks, Jackson) bearing B16-F10 melanoma tumors and receiving ICT (200 ug anti-PD-1 and 200 ug anti-CTLA-4 mAb).

**(B)** Cultured bacterial levels in MLN. MLN tissue homogenates were serially diluted, plated on BHI/Blood, YCFA, and CME0151 agar media and incubated at 37°C under anaerobic conditions for 24-72 hours. Quantification of colony-forming unit (CFU) from each agar plate was normalized to the tissue weight. n=7-8 per group. Points represent values from individual animals. Bars represent the mean ± SEM. Statistical analysis by Mann-Whitney test. *P<0.05, **P<0.01. ***, P<0.001.

**(C)** Relative abundance of cultured bacteria. Representative colonies (10-20 colonies with similar morphology and/or color) on each agar plate were selected and subsequently cultured in corresponding liquid media. gDNA was extracted. Full-length 16s rRNA gene (V1-V9 region) was amplified, purified, and sequenced (Sanger sequencing). Sequences were entered into the NCBI standard nucleotide Basic Local Alignment Search (BLAST) tool utilizing the rRNA/ITS databases. Bacterial species identification was ascertained from BLASTN results with the highest Total Score, with percent identity score >95% and E value <0.01.

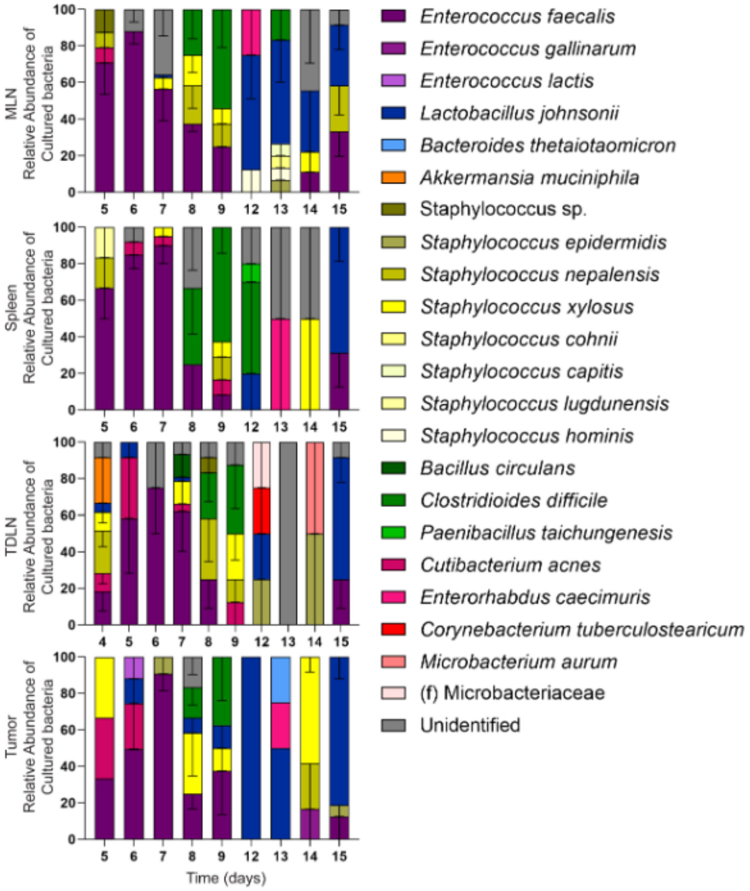


**Figure S2. Tissue and tumor microbiomes (cultured microbiota) in mice bearing B16-F10 tumors and receiving ICT, as described in fig 1A-C.**

Relative abundance of cultured bacteria from secondary lymphoid organ and tumor tissue recovered from C57BL/6J mice (n=6-8) bearing melanoma tumors and receiving anti-PD-1 and anti-CTLA-4 antibody treatment, as described in fig 1A. Tissue homogenates were serially diluted in reduced PBS and plated on YCFA agar and incubated for 24-72 hours at 37°C under anaerobic conditions. Colony-forming units (CFUs) were counted. Representative colonies based on morphology (~10-20 for each morphology type) were picked and cultured in YCFA medium for additional 24-48 hours at 37°C under anaerobic conditions. gDNA was isolated from bacterial cultures. Full-length 16s rRNA gene (V1-V9 region) was amplified, purified, and sequenced (Sanger sequencing). Sequences were entered into the NCBI standard nucleotide Basic Local Alignment Search (BLAST) tool utilizing the rRNA/ITS databases. Bacterial species identification was ascertained from BLASTN results with the highest Total Score, with percent identity score >95% and E value <0.01.

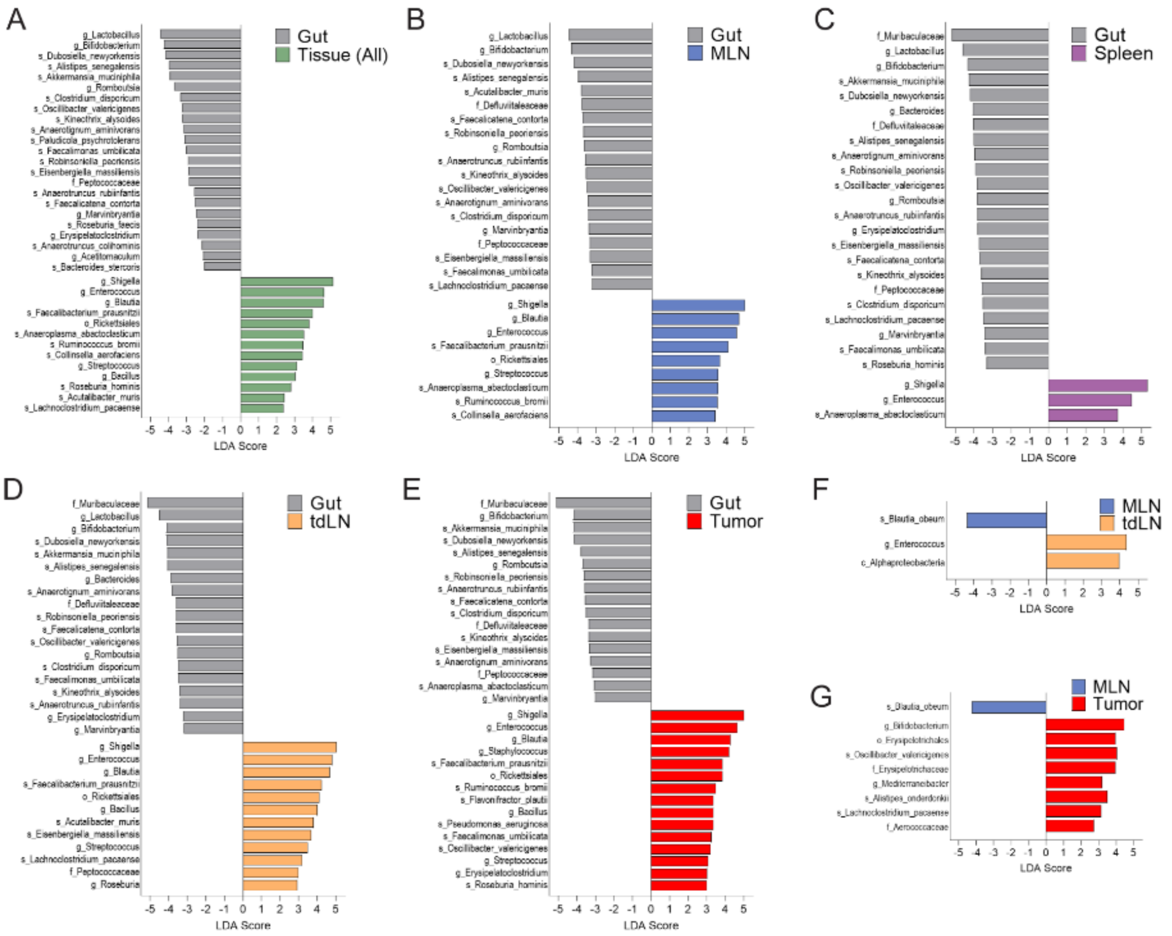


**Figure S3. Comparison of gut and tissue microbiomes from mice bearing melanoma tumors and receiving anti-PD-1 and anti-CTLA-4 antibody treatment.**
Microbiome composition determined by analysis of 16S rRNA sequencing (V4 region) of tissue, tumor and fecal samples recovered from C57BL/6J mice bearing melanoma tumors and receiving anti-PD-1 and anti-CTLA-4 antibody treatment, as in fig 1D-E. Differential bacterial taxonomic abundance between tissues types was analyzed by linear discriminant analysis effect size (LEfSe) projected as histograms. All listed bacterial groups (class (c), order (o), family (f), genus (g), or species(s)) were significantly enriched (>2 log-fold increase in linear discriminate analysis, LDA, score and $P<0.05$, Kruskal-Wallis test).

**(A)** Gut versus all other tissue types

**(B)** Gut versus mesenteric lymph node (MLN)

**(C)** Gut versus spleen,

**(D)** Gut versus tumor-draining lymph node (TDLN)

**(E)** Gut versus tumor

**(F)** MLN versus TDLN

**(G)** MLN versus tumor

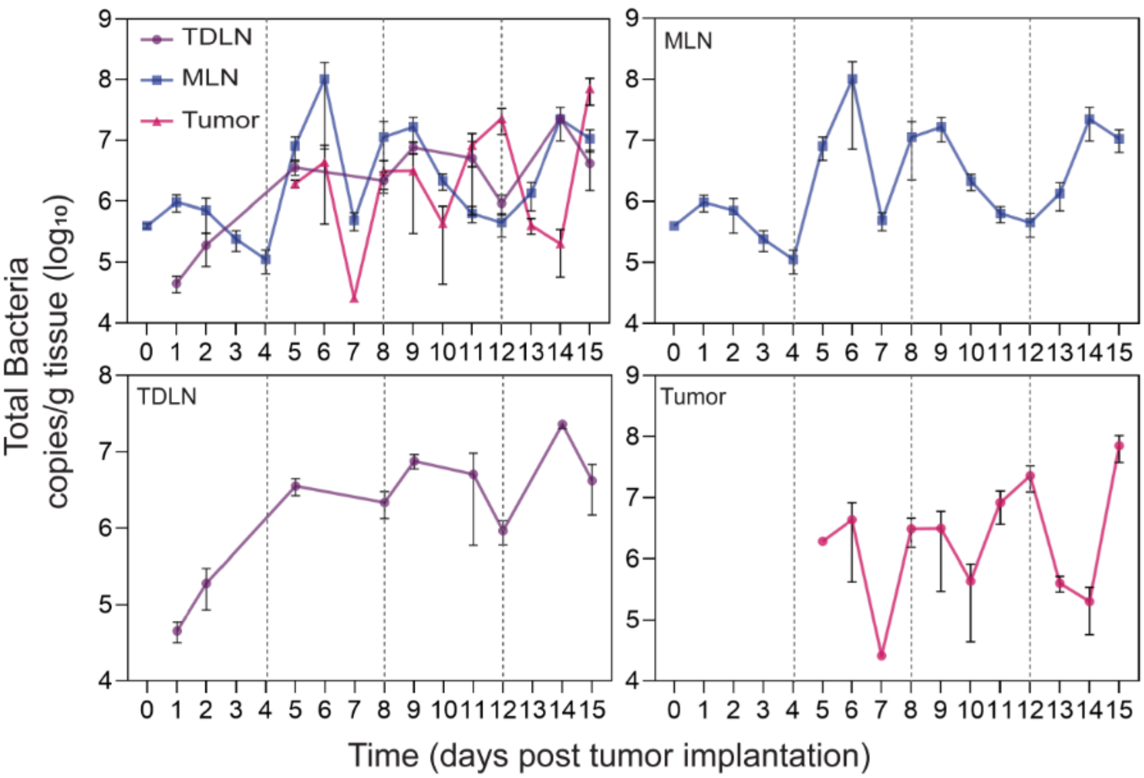


**Figure S4. Bacterial levels in MLN, TDLN, and tumor in mice bearing melanoma tumor and receiving ICT**

Bacterial load was quantified by qPCR-based quantification of 16s rRNA gene copies of tissue samples recovered from C57BL/6J mice (n=6-8) bearing melanoma tumors and receiving anti-PD-1 and anti-CTLA-4 antibody treatment, as described in fig 1A. Bacterial abundance determined by bacterial group (Eubacteria, all bacteria) quantitative-PCR (qPCR) of tissue gDNA collected from mice as detailed in fig. 1A.

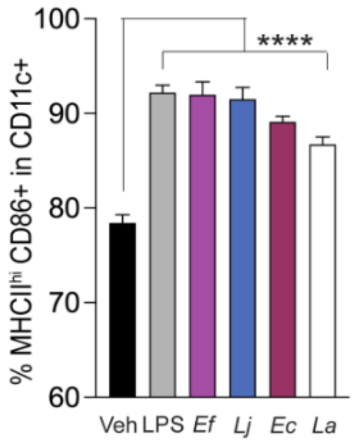


**Figure S5. CD86 expression in dendritic cells stimulated with different bacteria.** CD11c+ DCs were isolated from the spleen of C57BL6/J mice (female, 6-8 wks, Jackson) bearing B16-FLT3L tumors. Isolated DCs were stimulated with vehicle (PBS), *E. coli* LPS (Invivogen, 1ug/ml), and the following bacterial lysates (10ug/ml as determined by protein concentration via BCA assay): *Enterococcus faecalis* (*Ef*, clinical isolate from pediatric SCT patient), *Lactobacillus johnsonii* (*Lj*, VPI 7960), *Escherichia coli* (*Ec*, ATCC 10798), and *Lactobacillus acidophilus* (*La*, ATCC 4357) for 6 hours. Surface expression of MHC-II and CD86 were measured by flow cytometry. The proportion of MHC-II high, CD40+ cells among CD11c+ DCs are shown in the plot. Bars represent the mean ± SEM. Statistical analysis by Mann-Whitney test. ****$P<0.0001$. All assays were performed in triplicate.

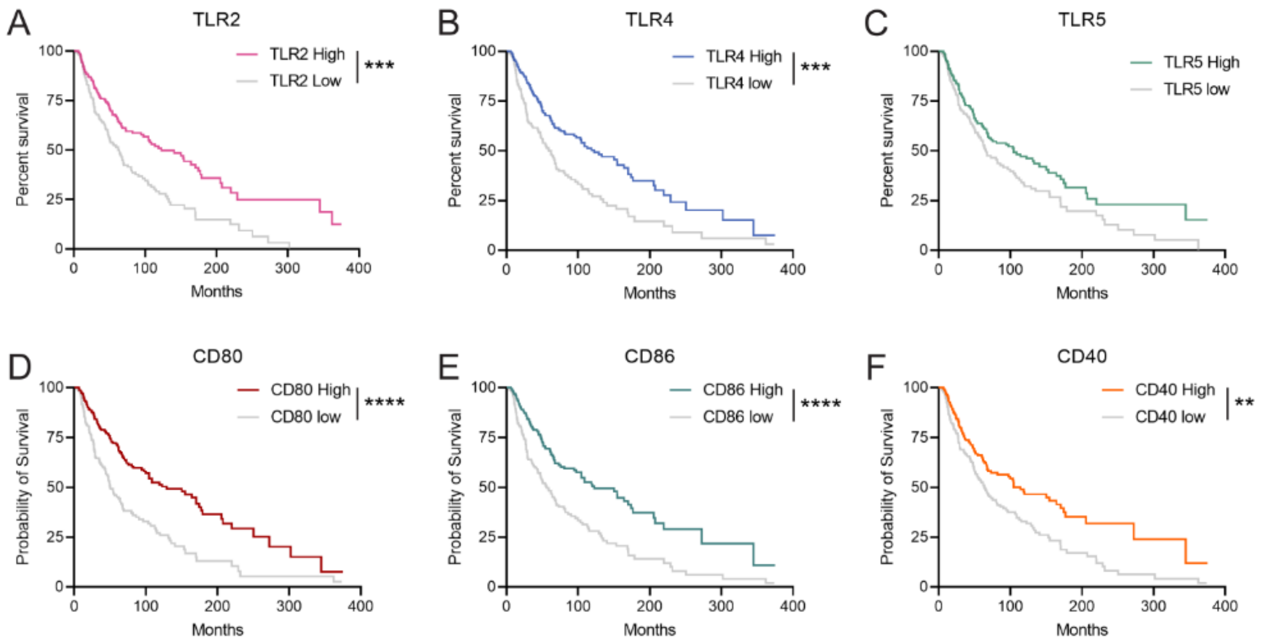


**Figure S6. Overall survival analysis of the Cancer Genome Atlas Human Skin Cutaneous Melanoma (TCGA-SKCM) cohort based on intratumoral mRNA expression of toll-like receptor (TLR) and co-stimulatory receptor genes.** The human skin cutaneous melanoma (SKCM) patient cohort (n=458) from The Cancer Genome Atlas (TCGA) was segmented into two groups, high (top 50%) and low (bottom 50%), based on the expression of TLR and co-stimulatory receptor genes. Kaplan-Meier plot of overall survival of the two patient segments based on the expression of (A) TLR2, (B) TLR4, (C) TLR5, (D) CD80, (E) CD86, and (F) CD40. Statistical analysis by Cox regression. *P<0.05, **P<0.01, ***P<0.0.001, ****P<0.0.0001.

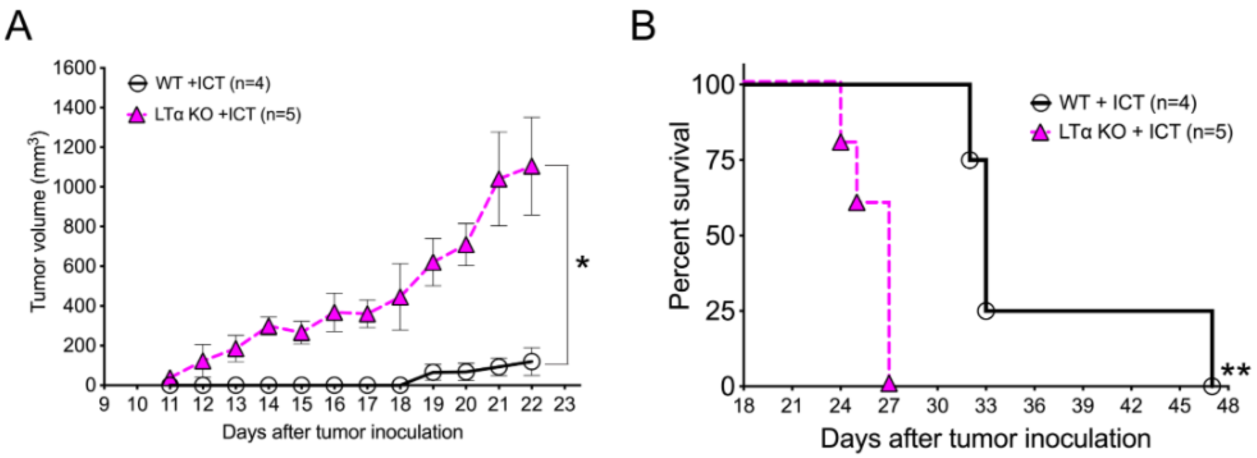


**Figure S7. Lymphotoxin alpha knockout (LTα KO) mice have diminished response to ICT compared to co-housed wild-type mice.**

(A) Tumor volumes (points equal mean ± SEM, Mann-Whitney test, *, $P < 0.05$) and (B) survival curves (log-rank test, **, $P < 0.01$) of lymphotoxin alpha knockout mice and wild-type mice (C57BL/6J, Jackson, female, 6-8, weeks old) implanted with B16-F10 tumors and treated with ICT (200 ug anti-PD-1 and 200 ug anti-CTLA-4 mAb) on days 4, 8 and 12 after tumor implantation.

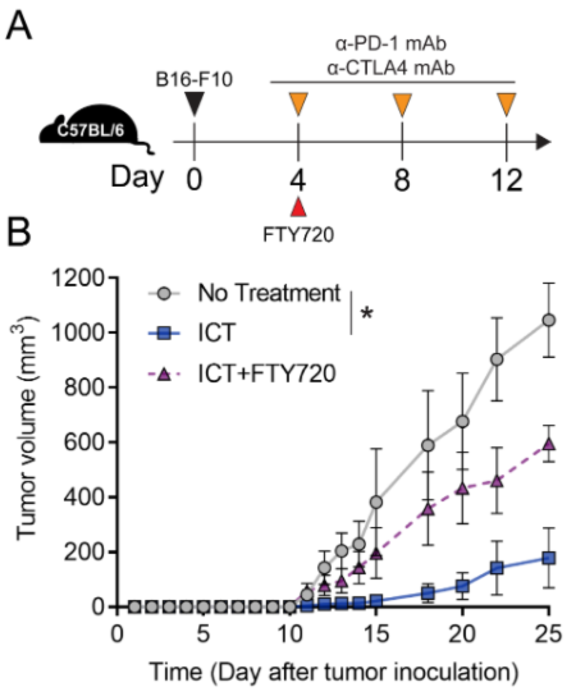


**Figure S8. FTY720-induced inhibition of lymphocyte egress from lymph nodes mitigates ICT efficacy.**

(**A**) Schematic overview of protocol. C57BL/6 mice (female, 6-8 wks, Jackson) were subcutaneously implanted with $1x10^5$ B16-F10 cells. Mice were injected intraperitoneally with ICT (200 µg anti-PD-1 and 200 µg anti-CTLA-4 mAb) on days 4, 8, and 12 post tumor inoculation. A single dose of FTY720 (Fingolimod, 0.3 mg/kg) was injected intraperitoneally on day 4 post tumor inoculation. n=4 per group.

(**B**) Tumor growth of mice treated ICT ± FTY720 or no treatment control. Points represent the mean ± SEM. Statistical analysis by Mann-Whitney test. *P<0.05.

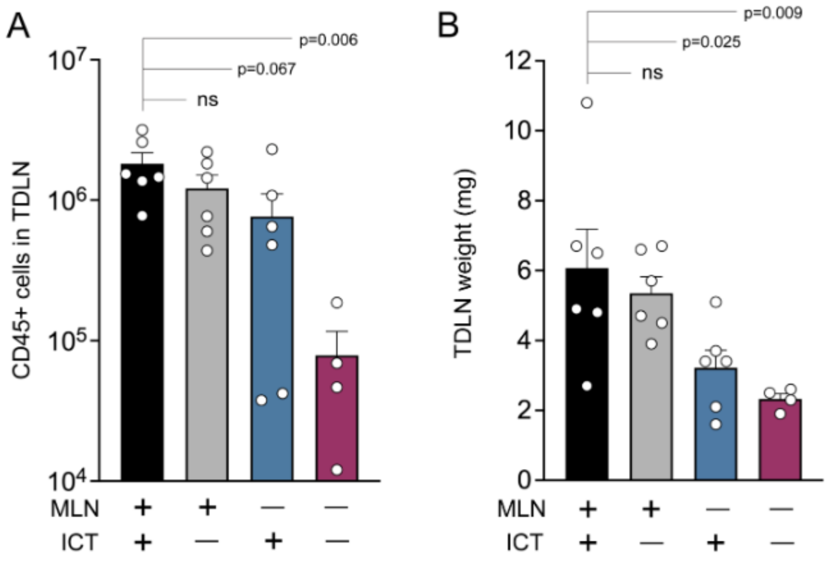


**Figure S9. Surgical resection of MLN results in decreased TDLN weight and lymphocytes.**

Tumor draining lymph nodes (TDLN, right inguinal lymph nodes) were harvested from mice (C57BL/6J, female, 6-8 weeks, n=4-6 per group) bearing B16-F10 tumors ± MLN ± ICT (anti-PD-1 and anti-CTLA-4 mAb) (as in fig. 3F). Isolated TDLNs were weighed and total TDLN CD45+ cell counts were measured by cell counting and flow cytometry.

**(A)** Quantification of CD45+ cells in the TDLN.

**(B)** TDLN weight.

Points represent values from individual animals. Bars represent the mean ± SEM. Statistical analysis by Mann-Whitney test.

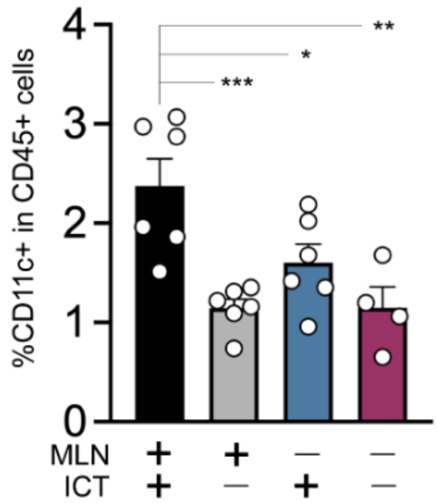


**Figure S10. Surgical resection of MLN results in decreased dendritic cells within TDLN of mice.**

The proportion of dendritic cells (CD11c+) among total tumor-infiltrating leukocytes (live CD45+) in TDLN of C57BL/6J mice (female, 6-8 weeks) with B16-F10 tumors ± MLN (via surgical resection) ± ICT (anti-PD-1 and anti-CTLA-4 mAb) (as in fig. 3F). The proportion of dendritic cells was analyzed by flow cytometry. n=4-6 per group.

Points represent values from individual mice. Bars represent the mean ± SEM. Statistical analysis by Mann-Whitney test. *P<0.05, **P<0.01, ***P<0.0.001.

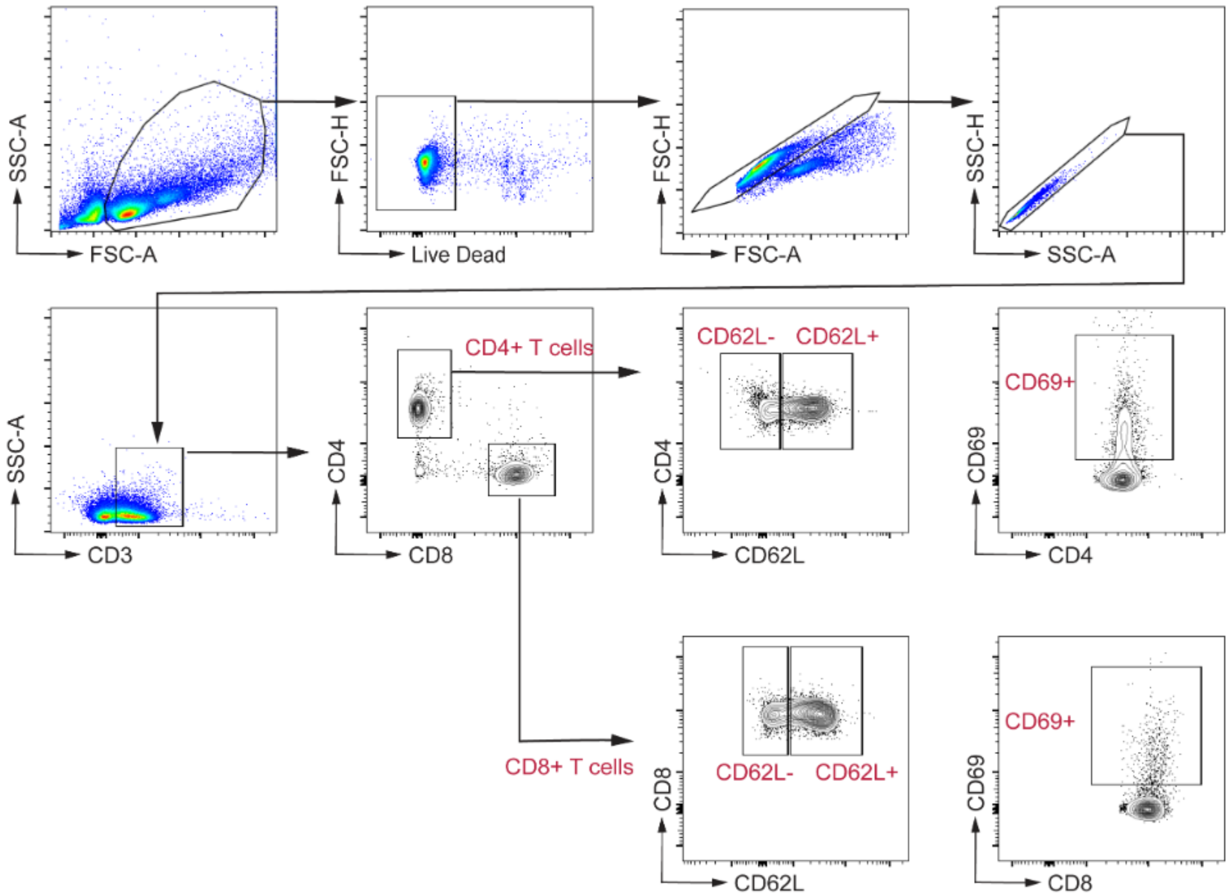


**Figure S11. Flow cytometry gating strategy for profiling TDLN T-cell phenotypes.**

Cells from the tumor-draining lymph nodes (right inguinal lymph nodes) were isolated and analyzed by flow cytometry. CD62L and CD69 expression within CD4+ T cells (live CD45+CD3+CD4+) and CD8+ T cells (live CD45+CD3+CD8+) were gated as shown. Representative plots are presented here and quantification of CD62L and CD69 expression on each cell population is shown in fig. 3 G-J.

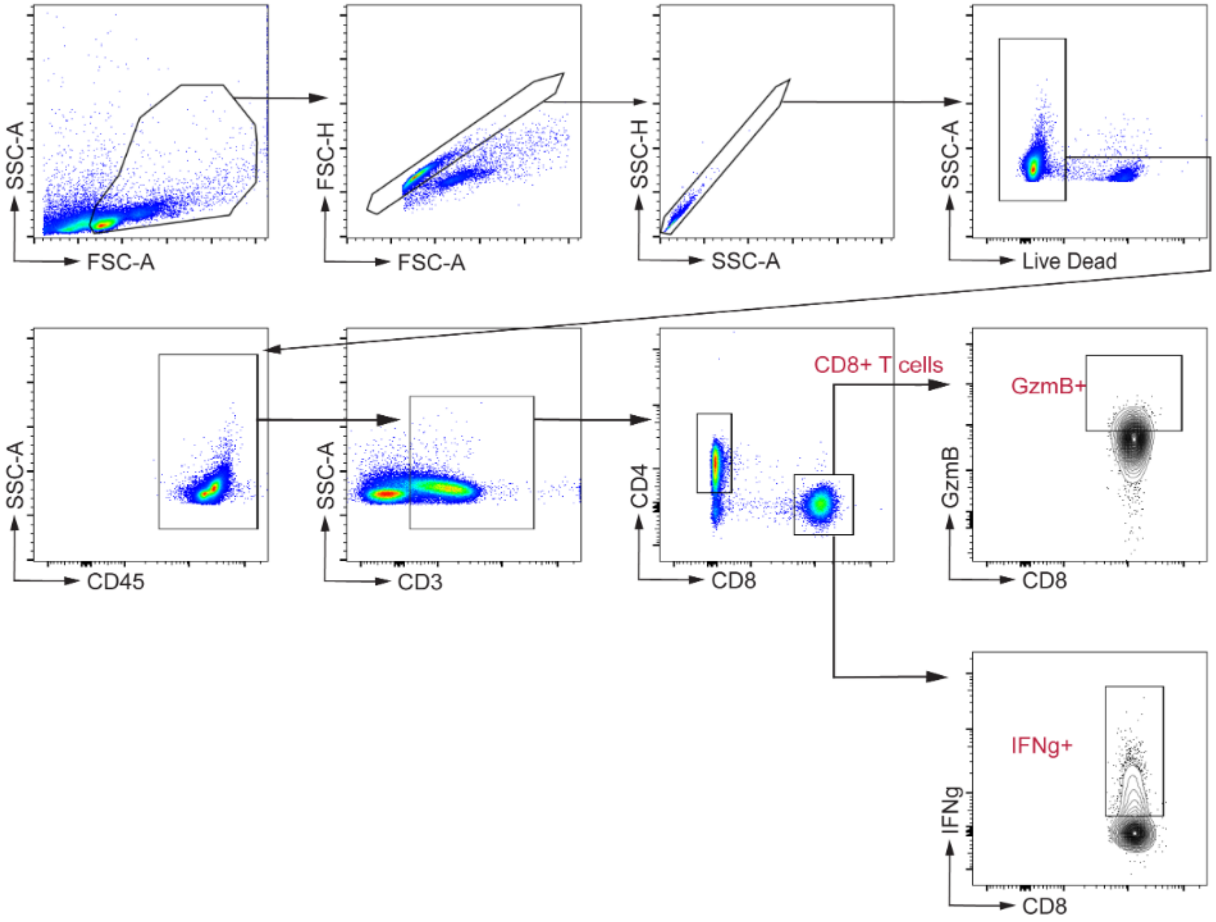


**Figure S12. Flow cytometry gating strategy for assessing TDLN T cell cytokine expression.** Cells from the tumor-draining lymph nodes were isolated and analyzed by flow cytometry. Intracellular expression of granzyme B (GzmB) and interferon-gamma (IFNg) within CD8+ T cells (live CD45+CD3+CD8+) were gated as shown. Representative plots are presented here and quantification of GzmB and IFN-γ expression on CD8+ T cell population is shown in fig. 3 K and L.

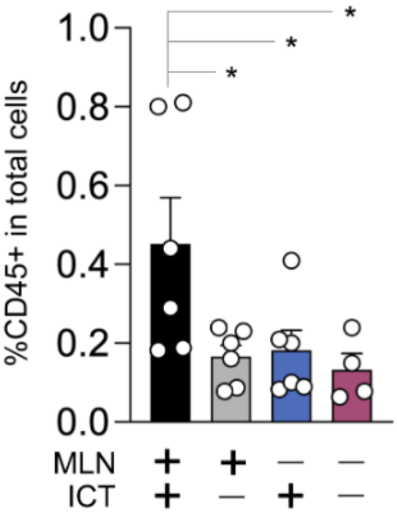


**Figure S13. MLN resection results in decreased leukocyte infiltrate in tumors of mice bearing melanoma tumors and receiving ICT.**

Tumor-infiltrating leukocytes were isolated from B16-F10 tumors from C57BL/6J mice (female, 6-8 weeks) ± MLN (via surgical resection) ± ICT (anti-PD-1 and anti-CTLA-4 mAb) (as in fig. 3F). The proportion of the tumor-infiltrating leukocytes (live CD45+) was analyzed by flow cytometry. n=4-6 per group. Bars represent the mean ± SEM. Statistical analysis by Mann-Whitney test. *P<0.05, **P<0.01, ***P<0.0.001.

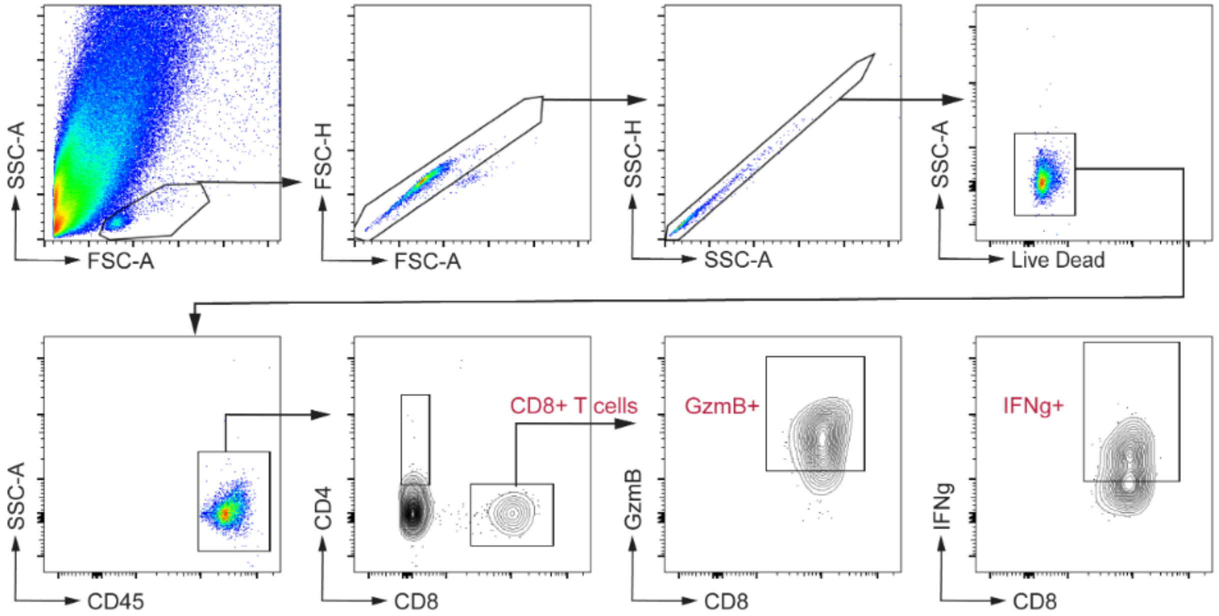


**Figure S14. Flow cytometry gating strategy for assessing tumor infiltrating leukocyte cytokine expression.**

Tumor-infiltrating leukocytes (CD45+) were isolated and analyzed by flow cytometry. Intracellular expression of granzyme B (GzmB) and interferon-gamma (IFNg) within CD8+ T cells (live CD45+CD8+) were gated as shown. Representative plots are presented here and quantification of GzmB and IFNg expression on CD8+ T cell population is shown in fig. 2 M and N.

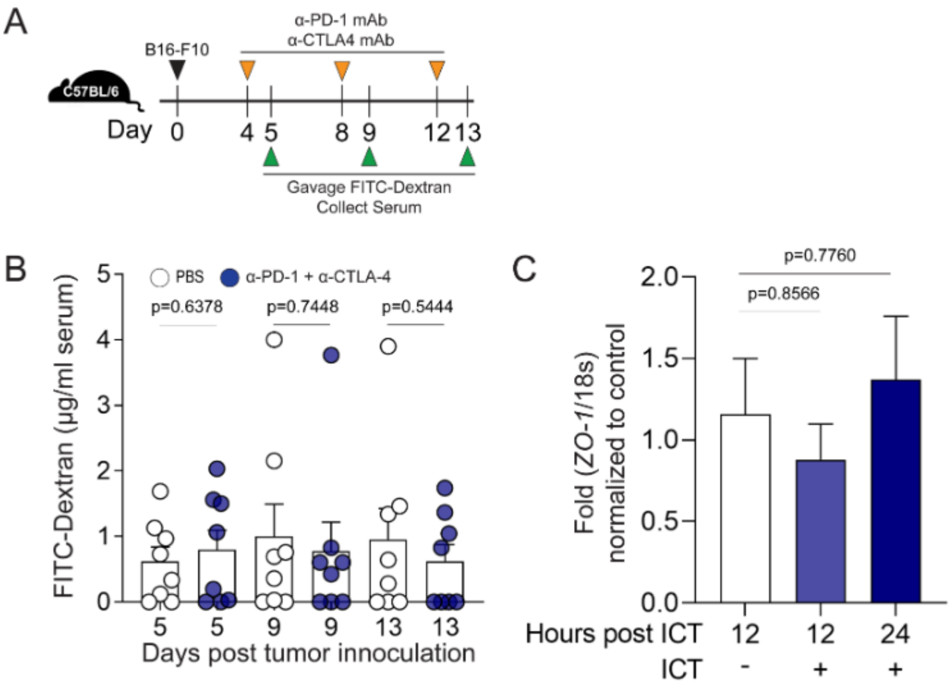


**Fig. S15. Gastrointestinal barrier function assays in mice bearing melanoma tumors and receiving anti-PD-1 and anti-CTLA-4 antibody therapy.**

(**A**) Schematic overview of protocol. C57BL/6 mice (female, 6-8 wks, Jackson) were subcutaneously implanted with $1x10^5$ B16-F10 cells. Mice were injected intraperitoneally with ICT (200μg anti-PD-1 and 200μg anti-CTLA-4 mAb) on days 4, 8, and 12 post tumor inoculation. Mice were fasted overnight and then orally gavaged with FITC-Dextran (4kD, 500mg/kg, Sigma). Serum was obtained 4 hours after gavage. Serum FITC-Dextran level was measured at an excitation wavelength of 485 nm and an emission wavelength of 528 nm.

(**B**) Serum FITC-Dextran levels in mice. n=7 per group. Points represent results from individual animals. Bars represent the mean $\pm$ SEM. Statistical analysis by unpaired t-test.

(**C**) Colonic expression of zonulin (*ZO-1*) in mice after receipt of ICT. Colons were collected 12 hours and 24 hours post ICT. mRNA expression of tight junction protein Zonula occludens-1, *ZO-1*, was measured by quantitative-PCR and normalized to 18s rRNA expression level. Bars represent the mean $\pm$ SEM. Statistical analysis by Anova test.

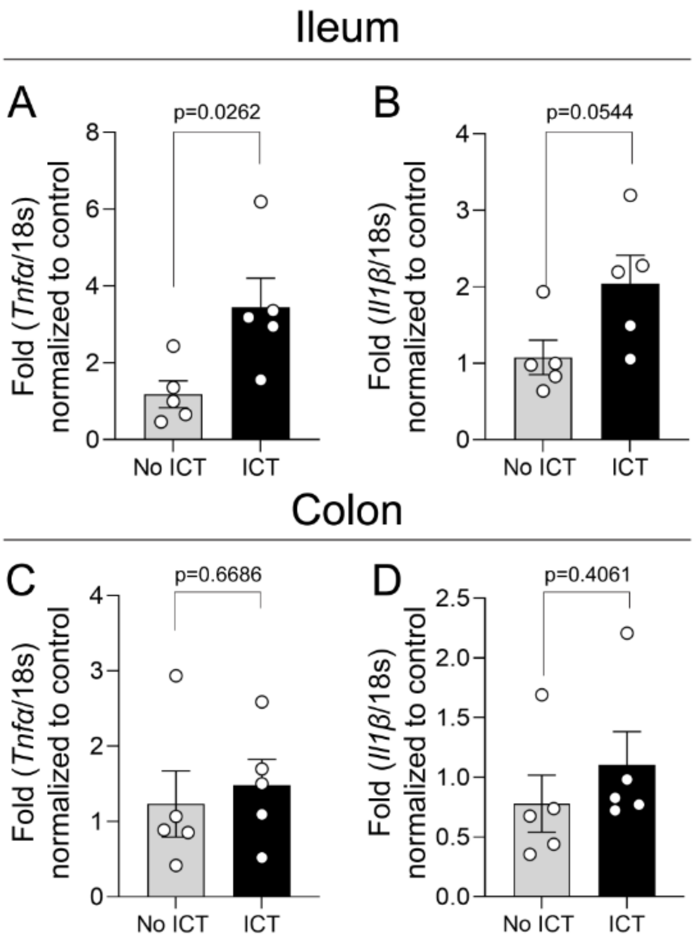


**Fig. S16. anti-PD-1 and anti-CTLA-4 therapy induces intestinal expression of TNF-*α* and IL-1β in mice.**

Ileum and colonic tissue was collected from C57BL6/J mice (Jackson, female, 6-8 weeks) bearing B16-F10 melanoma tumors and treated ± ICT (200 ug anti-PD-1 and anti-CTLA-4 mAb). Tissue samples were harvested one day after the first ICT dose (D+4 after tumor implantation).

(**A**) *Tnfa* and (**B**) *Il1b* mRNA expression in ileums from mice treated ± ICT

(**C**) *Tnfa* and (**D**) *Il1b* mRNA expression in colons from mice treated ± ICT

n=5 per group. Points represent values form individual animals. Bars represent the mean ± SEM. Statistical analysis by t-test. *P<0.05.

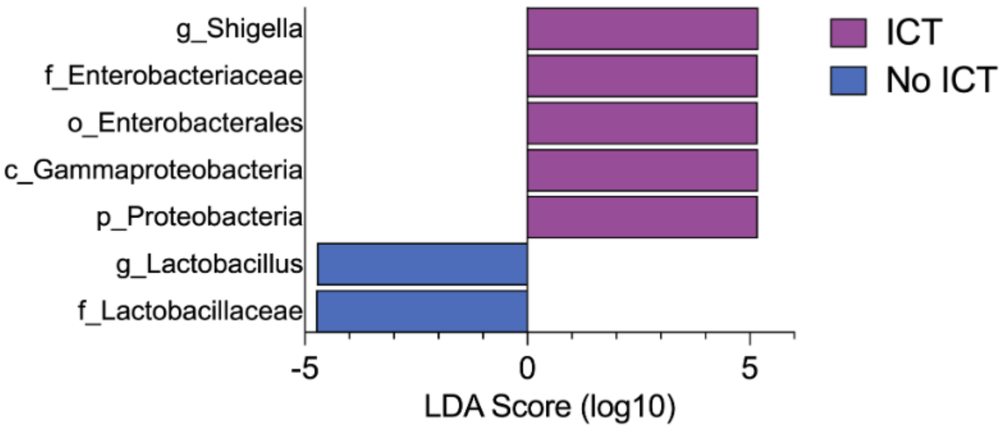


**Figure S17. Comparison of dendritic cell microbiomes from mice bearing melanoma tumors and treated with or without anti-PD-1 and anti-CTLA-4 antibody treatment.** Microbiome composition determined by analysis of 16S rRNA sequencing (V4 region) of dendritic cell samples recovered from MLNs in C57BL/6J mice bearing melanoma tumors treated with or without anti-PD-1 and anti-CTLA-4 antibody treatment (ICT). Differential bacterial taxonomic abundance between groups was analyzed by linear discriminant analysis effect size (LEfSe) projected as histograms. All listed bacterial groups (phylum (p), class (c), order (o), family (f), or genus (g)) were significantly enriched (>2 log-fold increase in linear discriminate analysis, LDA, score and $P<0.05$, Kruskal-Wallis test).

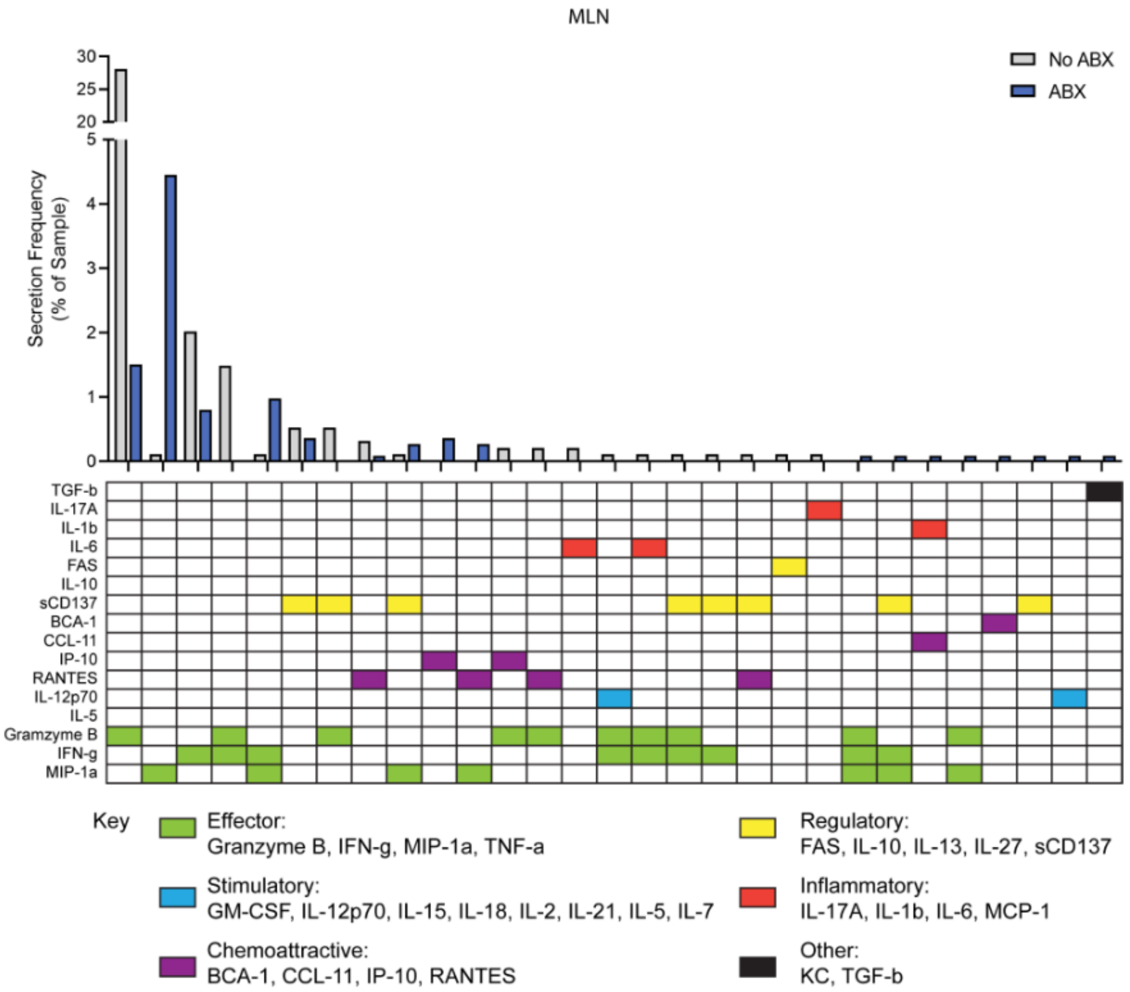


**Fig S18. Relative proportion of MLN CD8+ T cells isolated from mice + antibiotics (ABX) secreting different combinations of cytokines.**

C57BL/6 mice (female, 6-8 wks, Jackson) were treated + antibiotics (ABX, 2 mg/ml streptomycin and 1500 U/ml penicillin G in drinking water) for 7d before B16-F10 tumor inoculation. Mice were treated with 200μg anti-PD-1 and 200μg anti-CTLA-4 mAb intraperitoneally on days 4, 8, and 12 after tumor implantation. n=3 mice per group. Secretory cytokine profiles of CD8+ T-cells isolated from MLN of mice + ABX + ICT, as determined by single-cell multiplex cytokine profiling (Isoplexis IsoSpark; 28-plex mouse adaptive immune IsoCode chip panel).

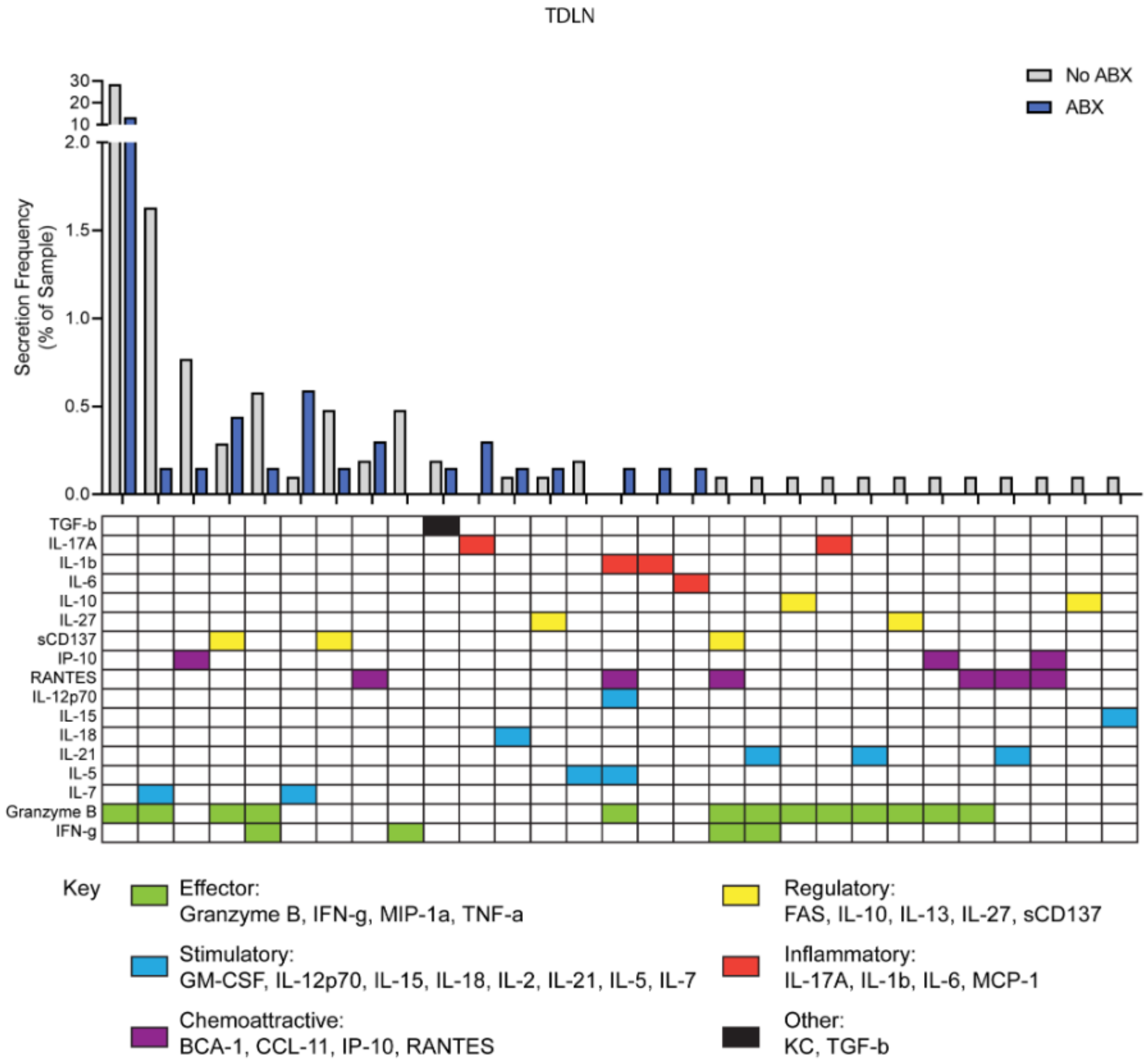


**Fig S19. Relative proportion of TDLN CD8+ T cells isolated from mice ± antibiotics (ABX) secreting different combinations of cytokines** C57BL/6 mice (female, 6-8 wks, Jackson) were treated ± antibiotics (ABX, 2 mg/ml streptomycin and 1500 U/ml penicillin G in drinking water) for 7d before B16-F10 tumor inoculation. Mice were treated with 200µg anti-PD-1 and 200µg anti-CTLA-4 mAb intraperitoneally on days 4, 8, and 12 after tumor implantation. n=10 mice per group. Secretory cytokine profiles of CD8+ T-cells isolated from TDLN of mice ± ABX + ICT, as determined by single-cell multiplex cytokine profiling (Isoplexis IsoSpark; 28-plex mouse adaptive immune IsoCode chip panel).

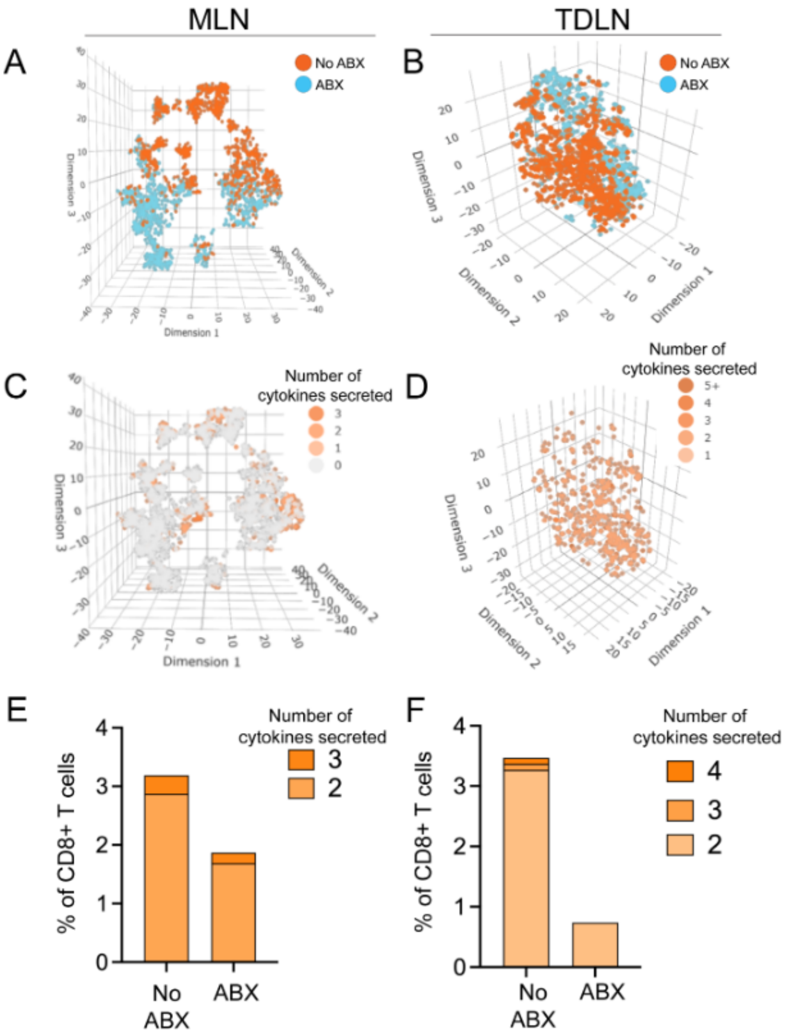


**Fig S20. Cytokine secretion profile of CD8+ T cells isolated from MLN and TDLN of mice treated with ICT and with or without antibiotics.**

Three-dimensional t-distributed stochastic neighbor embedding (t-SNE) plot of secretory cytokine profiles of CD8+ T-cells isolated from (**A**) MLN and (**B**) TDLN of B16-F10 melanoma tumor bearing mice (C57BL/6J, Jackson, female, 6-8 wks, n=3 for MLN group, n=10 for TDLN group) treated with ICT $\pm$ antibiotics (ABX, 2 mg/ml streptomycin and 1500 u/ml penicillin G). Representation of polyfunctional CD8 T cells from (**C**) MLN and (**D**) TDLN in the 3D-tSNE plot. The proportion of polyfunctional cells among total (**E**) MLN and (**F**) TDLN CD8+ T cells.

Supplementary Table 1: TCGA analysis raw statistics

| TCGA ID | TCGA Name | Updated ID | Updated Name | Cox coefficient | Raw p-value | BH-adjusted p-value | Median Expression | Mean Expression |
|---|---|---|---|---|---|---|---|---|
| 7097 | TLR2 | 7097 | TLR2 | -0.326 | 2.80E-06 | 2.56E-04 | 228.2908 | 344.0228688 |
| 7099 | TLR4 | 7099 | TLR4 | -0.266 | 1.20E-04 | 3.09E-03 | 233.0291 | 357.4088029 |
| 7100 | TLR5 | 7100 | TLR5 | -0.1015 | 1.50E-01 | 3.33E-01 | 46.7458 | 58.00507571 |
| 941 | CD80 | 941 | CD80 | -0.3445 | 2.90E-07 | 9.32E-05 | 11.6239 | 23.49641503 |
| 942 | CD86 | 942 | CD86 | -0.3292 | 8.40E-07 | 1.34E-04 | 160.3053 | 260.1028699 |
| 958 | CD40 | 958 | CD40 | -0.2809 | 6.80E-05 | 2.07E-03 | 254.5555 | 392.8295049 |

**Table S2. Antibodies used for flow cytometry**

Supplementary Table 2. Antibodies used for flow cytometry

| Specificity | Clone | Isotype | Fluorochrome | Supplier | Catalog number | Dilution |
|---|---|---|---|---|---|---|
| Live/Dead | | | Zombie yellow | Biolegend | 423104 | 1/600 |
| CD45 | 30-F11 | Rat IgG2b, κ | Brilliant violet 650 | Biolegend | 103151 | 1/200 |
| CD45 | 30-F11 | Rat IgG2b, κ | PE-Cyanine7 | Biolegend | 103114 | 1/200 |
| CD3e | 145-2C11 | Armenian Hamster IgG | APC | Biolegend | 100312 | 1/200 |
| CD4 | RM4-5 | Rat IgG2a, κ | AlexaFluor700 | Biolegend | 100536 | 1/200 |
| CD8a | 53-6.7 | Rat IgG2a, κ | FITC | Biolegend | 100706 | 1/200 |
| IFN-γ | XMG1.2 | Rat IgG1, κ | Pacific Blue | Biolegend | 505818 | 1/200 |
| Granzyme B | QA16A02 | Mouse IgG1, κ | PE-Cyanine7 | Biolegend | 372214 | 1/200 |
| CD62L | MEL-14 | Rat IgG2a, κ | Brilliant violet 510 | Biolegend | 104441 | 1/200 |
| CD69 | H1.2F3 | Armenian Hamster IgG | PE | Biolegend | 104508 | 1/200 |
| CD69 | H1.2F3 | Armenian Hamster IgG | Brilliant violet 650 | Biolegend | 104541 | 1/200 |
| CD11c | N418 | Armenian Hamster IgG | FITC | Biolegend | 117306 | 1/200 |
| MHC-II (I-A/I-E) | M5/114.15.2 | Rat IgG2b, κ | AlexaFluor700 | Biolegend | 107622 | 1/200 |
| CD80 | 16-10A1 | Armenian Hamster IgG | Brilliant violet 711 | Biolegend | 104743 | 1/200 |
| CD86 | GL-1 | Rat IgG2a, κ | PerCP | Biolegend | 105026 | 1/200 |
| CD40 | 3/23 | Rat IgG2a, κ | APC | Biolegend | 124612 | 1/200 |

### Table S3. Key resources table

| Table S3. Key resources table | | |
|---|---|---|
| **REAGENT or RESOURCE** | **SOURCE** | **IDENTIFIER** |
| **Antibodies (Other than flow cytometry)** | | |
| Anti-PD-1 (RMP-1, CD270) | BioXcell | Cat# BP0146 |
| Anti-CTLA-4 (9D9, CD152) | BioXcell | Cat# BP0164 |
| Anti-CD3e | Invitrogen | Cat# 16-0031-86 |
| Anti-CD28 | Invitrogen | Cat# 16-0281-85 |
| **Chemicals, peptides, and recombinant proteins** | | |
| FOXP3/Transcription Factor Staining Buffer kit | Invitrogen | Cat# 00-5521-00 |
| AccuPrime™ Pfx SuperMix | Invitrogen | Cat# 12344040 |
| SsoAdvanced Universal SYBR Green Supermix | Bio-Rad | Cat# 1725274 |
| Caprofen | Animal resource center, UTSW | DMC10-201013-01 |
| Recombinant murine IL-2 | Peprotech | Cat# 212-12 |
| Diphtheria Toxin from Corynebacterium diphtheriae | Sigma | Cat# D0564 |
| Ultrapure LPS, E. coli 0111:B4 | Invitrogen | Cat# tlrl-3pelps |
| **Experimental models: Organisms/strains** | | |
| Enterococcus faecalis | This study | |
| Lactobacillus johnsonii VPI 7960 | ATCC | ATCC 33200 |
| Lactobacillus acidophilus ATCC 4357 | ATCC | ATCC 4357 |
| Escherichia coli ATCC 10798 | ATCC | ATCC 10798 |
| **Experimental models: Cell lines** | | |
| B16-F10 | ATCC | ATCC CRL-6475; RRID:CVCL_0159 |
| B16-FLT3L | Dr. Chandrashekhar Pasare | RRID:CVCL_IJ12 |
| **Critical commercial assays** | | |
| Pierece BCA assay kit | Thermo Scientific | REF 23225 |
| CD11c microbeads ultrapure, mouse | Miltenyi Biotec | Cat# 130-125-835 |
| **Software and algorithms** | | |
| FlowJo (v.10.8.0) | BD Biosciences | https://www.flowjo.com/ |
| QIIME 2 | Bolyen et al., 2019 | https://qiime2.org |
| GraphPad Prism 9 | GraphPad Software | https://www.graphpad.com |
| OncoLnc | J Anaya, 2016 | http://www.oncolnc.org/ |
| **Other** | | |
| Screw-cap microfuge tube (2ml) | Fisher Scientific | REF 72.693.005 |
| Borosilicate glass beads (5mm) | Sigma | Cat# Z143944 |
| Pack-Rectangular Jar (2.5L) | Mitsubishi Gas Chemical | Order# 50-25 |
| Anaerobic gas pack | Mitsubishi Gas Chemical | Order# 10-01 |
| Absorbable sutures (5/0 PGA) | Covetrus | Cat# 031995 |
| Nonabsorbable sutures (5/0 Monofilament nlylon) | Covetrus | Cat# 056919 |
| Vetrinary surgical adhesive | Covetrus | Cat# 031477 |
| Triple antibiotic ointment (Bacitracin zinc, neomycin sulfate, and polymyxin b sulfate ointment | Taro Pharmaceutical | NDC 51672-2120-2 |
| Sterile cell strainer (70µm) | Fisher Scientific | Cat# 22363548 |
| Syringe (10ml) | BD | REF 302995 |
| Petridish | Falcon | Cat# 351029 |
| gentleMACS™ Dissociator | Miltenyi Biotec | Cat# 130-093-235 |
| Tumor dissociation kit, mouse | Miltenyi Biotec | Cat# 130-096-730 |
| LS column | Miltenyi Biotec | Cat# 130-042-401 |
| BD Vacutainer SST tube | BD | REF 367983 |
